# Immune modulation of innate and adaptive responses restores immune surveillance and establishes anti-tumor immunological memory

**DOI:** 10.1101/2023.09.27.559828

**Authors:** Ayesha B. Alvero, Alexandra Fox, Bhaskara Madina, Marie Krady, Radhika Gogoi, Hussein Chehade, Valerian Nakaar, Bijan Almassian, Timur Yarovinsky, Thomas Rutherford, Gil Mor

## Abstract

Current immunotherapies have proven effective in strengthening anti-tumor immune responses but constant opposing signals from tumor cells and surrounding microenvironment eventually lead to immune escape. We hypothesize that in situ release of antigens and regulation of both the innate and adaptive arms of the immune system will provide a robust and long-term anti-tumor effect by creating immunological memory against the tumor. To achieve this, we developed CARG-2020, a virus-like-vesicle (VLV). It is a genetically modified and self-amplifying RNA with oncolytic capacity and encodes immune regulatory genes. CARG-2020 carries three transgenes: 1) the pleiotropic antitumor cytokine IL-12 in which the subunits (p35 and p40) are tethered together; 2) the extracellular domain (ECD) of the pro-tumor IL-17RA, which can serve as a dominant negative antagonist; and 3) shRNA for PD-L1. Using a mouse model of ovarian cancer, we demonstrate the oncolytic effect and immune modulatory capacities of CARG-2020. By enhancing IL-12 and blocking IL-17 and PD-L1, CARG-2020 successfully reactivates immune surveillance by promoting M1 instead of M2 macrophage differentiation, inhibiting MDSC expansion, and establishing a potent CD8+ T cell mediated anti-tumoral response. Furthermore, we demonstrate that this therapeutic approach provides tumor-specific and long-term protection preventing the establishment of new tumors. Our results provide rationale for the further development of this platform as a therapeutic modality for ovarian cancer patients to enhance the anti-tumor response and to prevent recurrence.

## Introduction

The development of solid tumors requires modulation of the host immune system and successful escape from immune surveillance (1,2). This is accomplished by mechanisms that recruit and educate immune cells to support tumor growth and promote metastatic expansion (3-5). Immunotherapy, specifically immune checkpoint blockers, has transformed the treatment of several solid tumors and has demonstrated success in the modulation of the tumor microenvironment (TME), promotion of anti-tumor responses and consequently regression of tumors (6-8). Unfortunately, not all types of tumors have benefitted from this modality and currently available immune modulators have shown limited success with recurrent disease (9,10).

Conventional treatment for solid tumors such as ovarian cancer involves chemotherapy, which is administered prior or after surgery(11). Although chemotherapy eradicates majority of the tumor, undetectable micro-metastatic niches always persist (12). These chemoresistant cells are thought to drive relapse and in addition are known to create a unique microenvironment, which modulates both the local and systemic immune system. Therefore, immune therapies that can successfully target these chemoresistant micro-metastases could increase the likelihood of permanent remission.

One of the well-known mechanisms that confer resistance to immune therapies is the establishment of a “cold” TME (13,14). This is a state wherein cytolytic T cells are actively prevented from infiltrating the tumor and is attained by the recruitment of immune suppressor cells such as myeloid-derived suppressor cells from the bone marrow (BM-MDSC), M2 macrophages, and T regulatory (Treg) cells (15). In contrast, “hot” tumors are characterized by presence of T cells, antigen presenting cells (M1 macrophages and dendritic cells), and absence of immune suppressor cells consequently demonstrating better response to immune therapy and ultimately better outcomes (16). A conceivable path to improve T cell antitumor activity is to counteract immunosuppressive signals, such as those of BM-MDSC. Indeed, studies have shown that poor clinical outcomes are associated with an increase in tumor infiltration of immunosuppressive cells such as tumor-associated macrophages (TAMs) and BM-MDSC.

Another important characteristic of “cold” tumors is poor antigenicity, which further enhances the tolerogenic state of its microenvironment and impedes the success of anti-tumor vaccines. *In situ* vaccination (ISV) has been proposed as an efficient approach to remodel the tumor from “cold” to “hot” while boosting anti-tumor T cell response (17). However, the selection of the most efficient antigen, which represents the heterogeneity of the tumor has proven to be a major challenge (18-20). A promising strategy is to develop personalized vaccination by inducing cancer cell death and enhancing antigen presentation.

The use of oncolytic virus has been proposed as an effective mechanism to induce the release of tumor antigens as a result of virus-induced cell death (19,21,22). Oncolytic viruses infect/replicate only in cancer cells leading to cancer cell lysis. Its efficacy is amplified when used as an ISV by promoting cancer cells apoptosis, releasing high levels of tumor antigens, and provoking an anti-tumor immune response (23,24). However, although the oncolytic effect can enhance the release of tumor antigens, it is inefficient in modulating the components of immune system (23). It is plausible that the development of a more comprehensive immunotherapy strategy that engages multiple arms of the immune system and incorporates immune modulatory genes with the oncolytic effect can induce a more potent and persistent anti-tumor effect.

The objective of the study is to elucidate the efficacy of a novel therapeutic approach, CARG-2020, that can modulate immune responses while inducing cancer cells death. CARG-2020 is an armed oncolytic virus-like vesicle (VLV) vector. VLV is a hybrid vector with components from two unrelated animal viruses, the alphavirus Semliki Forest virus (SFV) and rhabdovirus vesicular stomatitis virus glycoprotein (VSV-G) and produces infectious replication-competent enveloped vesicles at high titers in vitro (25,26). VLV are a capsid-free, self-replicating virus-like vaccine platform carrying positive-strand capped and polyadenylated RNA encoding an *in vitro* evolved SFV RNA-dependent RNA replicase and the VSV glycoprotein. VLV can be engineered to express foreign antigens, proteins, or microRNAs (miRNA) that can modulate the immune response (27,28). The VLV platform replicates like a virus, but its only structural protein is the VSV glycoprotein (VSV-G), and unlike many other viral vectors, lacks pathogenicity (29,30). CARG-2020 is a membrane encapsulated oncolytic VLV RNA replicon delivering three immunomodulators: IL-12, dn-IL17RA (dominant-negative IL-17 receptor A), and shRNA for PD-L1.

Previously, we evaluated a VLV and its potential impact on cancer cells and observed that it can function in the context of oncolytic virus therapy (24). In this present study, we tested the capacity of a VSV-based oncolytic virus, CARG-2020, to function as an ISV by inducing cancer cell death and to modulate both the innate and adaptive arms of the immune system through the expression of three immune modulatory molecules: IL-12, IL-17R ECD (extra-cellular domain), and short hairpin RNA (shRNA) for PD-L1. Using human and mouse ovarian cancer models in both immunocompromised (human ovarian cancer cells) and immunocompetent mice (mice ovarian cancer cells), we demonstrate that CARG-2020 has potent oncolytic effect in cancer cells but not in normal cells and is well tolerated in vivo. In addition, we demonstrate that CARG-2020 can convert the “cold” ovarian TME to “hot” and provide specific long -term anti-tumor immunity capable of preventing tumor expansion and disease recurrence.

## Methods and methods

### Cell lines and culture conditions

TKO mouse ovarian cancer cells were kindly provided by Dr. Martin Matzuk. mCherry fluorescence was stably expressed using a lentivirus to allow monitoring of i.p. tumors in real time. TKO cells were cultured in DMEM/F12 media supplemented with 10% FBS and 1% Penicillin-Streptomycin.

OCSC1-F2 were isolated from tumors derived from ovarian cancer patients as previously described **(24,31-36)**. and cultured in RPMI media supplemented with 10% FBS, 1% Penicillin-Streptomycin, 1% Sodium Pyruvate, 1% HEPES, and 1% non-essential amino acids. Both cell lines were grown at 37°C with 5% CO_2._ Cells were routinely tested for mycoplasma and authenticated once a year by STR profiling and used within 6 passages between experiments.

### Generation of recombinant CARG-2020 vector and Particles

The VLV double promoter (dp) vector was for cloning was prepared by linearizing it with Asc I and Sbf I restriction enzymes in order to insert VSV G glycoprotein fragment from New Jersey (NJ) serotype downstream of the second sub-genomic promoter (27). The resulting VLVdp VSV G^NJ^ vector was then digested with BamH I and Pac I enzymes and the resulting vector was used to insert cytokines and shRNA downstream of the first sub-genomic promoter. In order to generate the CARG-2020 construct containing three transgenes namely IL-12, IL-17RA (Accession # NP_032385.1) and shRNA target sequences derived from PD-L1 (Accession # NM_021893.3), the full-length IL-12 fragment (encompassing both fused subunits), and an IL-17RA extracellular domain (ECD) containing a 3’ HA-tag and PD-L1-shRNA synthetic gene fragment incorporating three shRNA concatemers (Sh-1: 5’ATTTGCTGGCATTATATTCAC-3’; Sh-2: 5’-GCTGAAAGTCAATGCCCC ATA-3’ and Sh-3: 5’-CTGGACAAACAGTGACCACCA-3’) were amplified and infused into VLV dp - VSV G^NJ^ between BamHI/PacI cloning sites. To clone GFP into the VLV vector, an insert was amplified and cloned in VLV dp - VSV G^NJ^. All primers to amplify DNA fragments were designed according to NEBuilder instruction and infused into VLV dp - VSV G^NJ^ vector using NEB HiFi DNA Assembly Kit (E5520S). The recombinant clones were screened for positive inserts by DNA restriction digestion. The final positive constructs were further confirmed by DNA sequencing (GENEWIZ, NJ, USA).

The full-length IL-12 fragment, and an IL-17RA ECD HA tag/PD-L1-shRNA fragment driven by sub-genomic promoter sequence were amplified by PCR and infused into VLV dp VSV-G^NJ^ vector between BamHI/PacI cloning sites downstream of the first sub-genomic promoter. All primers to amplify DNA fragments were designed according to NEBuilder instruction and infused into VLV backbone using NEB HiFi DNA Assembly Kit (E5520S). The recombinant clones were screened for positive inserts by DNA restriction digestion. The final positive constructs were further confirmed by DNA sequencing (GENEWIZ, NJ). For the production of VLV stocks, BHK-21 cells were cultured in Dulbecco’s Modified Eagle Medium supplemented with 5% fetal bovine serum (FBS). VLVs were produced by transfecting BHK-21 cells with the VLV plasmid DNA followed by collection of the master VLV stock. Propagation of the working stocks was performed by a single passage of the master stock in BHK-21 cells cultured in Opti-MEM™ Reduced Serum Medium (ThermoFisher, Waltham, MA, USA). Working stocks were concentrated using MacroSep® Advance 100K MWCO (Pall Laboratories, Port Washington, NY, USA) and titrated using plaque assay in BHK-21 cells.

### Immunocompetent mouse model and treatment schedule

All the described experiments using mice were approved by Wayne State University Animal Care and Use Committee (IACUC 22-03-4474) and mice were housed at Wayne State University Division of Laboratory Animal Resources. 1×10^7^ mCherry^+^ TKO mouse ovarian cancer cells were injected intra-peritoneally (i.p.) in 6-8 week old female C57BL/6 mice. Tumor growth was monitored every 4 days by detecting mCherry fluorescence using Ami HT Imaging System (Spectral Instruments, Tucson, AZ). Tumor burden was quantified using mCherry fluorescent ROI area using Aura Imaging Software (Spectral Instruments). Treatment commenced when mCherry ROI area reached 3.2 x 10^8^ PFU/sec (around day 18 post injection of cells and at the beginning of the Expansion phase; Fig. 1A). CARG-2020 was administered i.p. at 1×10^6^ PFU/dose given every 3 days for a total of 3 doses. Animal weight and abdominal width were monitored twice a week. Animals were sacrificed when mCherry ROI area reach 1 x 10^9^ PFU/sec or when abdominal width reaches or exceeds 3.4 cm.

**Figure 1.**
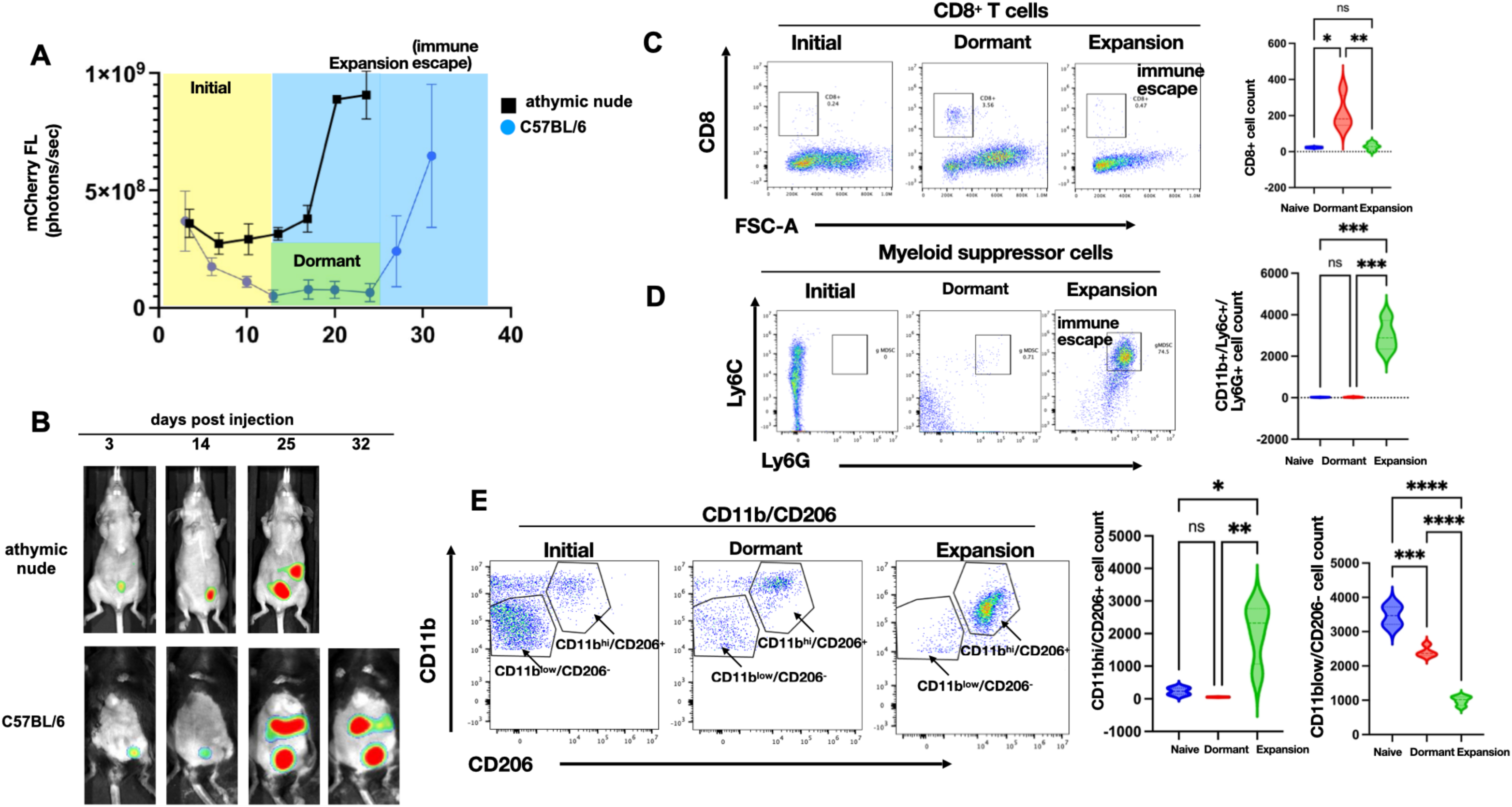
Characterization of immune response during ovarian tumor progression. mCherry+ TKO mouse ovarian cancer cells were injected i.p. in C57BL/6 immunocompetent mice (n=50) or immunocompromised athymic nude mice (n=10). **A.** Tumor growth was quantified using mCherry ROI fluorescence. Note difference in tumor kinetics between mouse strain with the dormant phase observed only in C57BL/6 mice; **B.** Representative images of live animal imaging showing progression of i.p. tumor burden; **C.** Peritoneal lavage from TKO-bearing C57BL/6 mice from each phase of tumor progression (n=6) was stained with anti-CD8 and analyzed by flow cytometry; **D.** Peritoneal lavage from TKO-bearing C57BL/6 mice from each phase of tumor progression (n=6) was stained with anti-CD11b, anti-Ly6C, and anti-Ly6G and analyzed by flow cytometry; **E.** Peritoneal lavage from TKO-bearing C57BL/6 mice from each phase of tumor progression (n=6) was stained with anti-CD11b and CD206 and analyzed by flow cytometry. Representative dot plots are shown. Gating strategy is shown in Supp. Fig. 1. Data are presented as Mean ± SEM and One-way ANOVA was used for statistical analysis.

### Immune-incompetent model and treatment schedule

5×10^6^ mCherry^+^ TKO mouse ovarian cancer cells were injected i.p. in 6-8 week old female athymic nide mice. Tumor growth was monitored and quantified as above. Treatment commenced when mCherry ROI area reached 3.2 x 10^8^ PFU/sec (around day 11 post injection of cells). CARG-2020 was administered i.p. at 1×10^7^ PFU/dose given every 3 days for a total of 3 doses. Animal weight and abdominal width were monitored twice a week. Animals were sacrificed when mCherry ROI area reach 1 x 10^9^ PFU/sec or when abdominal width reaches or exceeds 3.5 cm.

### Immunephenotyping by FACS analysis

Peritoneal cells were collected from ascites-free mice by i.p. injection of sterile PBS followed by fluid aspiration into heparin-containing tubes. Peritoneal fluid from ascites-containing mice was collected by peritoneal tap into heparin-containing tubes. Splenocytes were obtained by homogenizing spleens. After washing and RBC lysis, the following antibodies were used: APC-conjugated CD8, Superbright 785-conjugated Cd11b, APCFire 750-conjugated Ly6C, Pacific blue-conjugated Ly6G, APC-conjugated CD206, Brilliant violet 605-conjugated MHC-II, Superbright 785-conjugated CD44, FITC-conjugated CD62L. Antibodies were purchased from ThermoFisher and Biolegend. Flow cytometry was carried out on with electronic gates set on live and single cells. A minimum of 5 × 10^4^ events was collected per sample, and data were analyzed with FlowJo software (FlowJo LLC). Gating strategy are shown in Supplementary Figure 1.

### RNA extraction, cDNA synthesis and qPCR

Total RNA was extracted from freshly isolated cell pellets using RNAeasy Mini Kit (Qiagen, Austin, TX) according to the manufacturer’s instructions and with DNase treatment. RNA was quantified and purity was assessed using Epoch Microplate Spectrophotometer (Agilent, Santa Clara, CA). cDNA was synthesized using iScript cDNA kit (Bio-Rad, Hercules, CA) from 1 microgram of total RNA according to manufacturer’s instructions. qPCR was performed using SYBR Green Supermix (Bio-Rad, Hercules, CA) with 1:10 dilution of cDNA in a final volume of 10 μl according to manufacturer’s instructions. qPCR was run on CFX96TM PCR detection system (Bio-Rad, Hercules, CA) using the following thermocycling parameters: initial denaturation step of 2 minutes at 95°C; Primer annealing step for 30 seconds at 55°C and elongation step for 60 seconds at 74°C for 30 cycles; and final extension step for 5 minutes at 74°C. Primers for IL-12 were synthesized by Integrated DNA Technologies (San Diego, CA) and sequences were as follows. IL-12: forward, 5’-CACACTGGACCAAAGGGAC-3’; reverse, 5’-CAAAGGCTTCATCTGCAA-3’. Ppia was used a reference gene: forward, 5’-AGCACTGGGGAGAAAGGATT-3’; reverse, 5’-AGCCACTCAGTCTTGGCAGT-3’. Control group was treated with PBS. No RT control was used as negative control. Relative expression was calculated using the comparative ΔΔCT method with Control group as reference. All reactions were performed in triplicates.

### Western blot analysis

Whole cell protein lysates were isolated by resuspending cell pellets in 1X Cell lysis buffer (Cell Signaling Techonologies, Danvers, MA) with added cOmplete^TM^ Protease Inhibitor Cocktail (Millipore Sigma, Burlington, MA) and centrifugation for 20 minutes at 1,500 rpm. 20 mg of protein lysate was electrophoresed on 12% SDS-polyacrylamide gels then transferred to PVDF membranes (EMD Millipore, Burlington, MA). After blocking with 5% Milk, membranes were probed with primary antibodies at 4°C overnight, and then secondary antibodies for 1 hour at room temperature. The blots were developed using enhanced chemiluminescence and imaged using GE ImageQuant LAS 500 chemiluminescence (Cytiva Life Sciences, Marlborough, MA).

### Cell growth and cell death assays

3,000 TKO cells were seeded per well of a 96-well plate and allowed to reach ∼80% confluency. Increasing dose of CARG-2020 was added in a 200 ul total volume with added Celltox^TM^ fluorescent dye (Promega, Madison, WI) at 1;2,000 final dilution. Plates were placed in Cytation5/Biospa (Agilent Technologies, Santa Clara, CA) for automatic imaging. Images were analyzed using Gen5 (Agilent Technologies).

### Splenocyte transfer

Splenocytes were triturated from spleens and cells were collected in PBS and passed through a 100 um mesh filter. Cells were counted after washing and RBC lysis. 5×10^5^ splenocytes were injected i.p. to recipient mice.

### Statistical analysis

Unpaired two-tailed Student t-tests assuming Gaussian distribution or one way analysis of variance (ANOVA) with Dunnett’s multiple comparisons were used for comparison between different groups. P values of 0.05 or less were considered statistically significant. Statistical analysis was performed and all data were graphed using GraphPad Prism v9.3.1(San Diego, CA; RRID:SCR_002798). Data are presented as mean ±SEM.

### Data availability statement

Data sharing is not applicable to this article as no data were created or analyzed in this study.

## Results

### Carcinomatosis is associated with an immunosuppressive peritoneal microenvironment

To characterize the temporal regulation of the immune response during the process of carcinomatosis formation, we utilized a previously reported syngeneic mouse model of ovarian cancer established using mCherry-expressing (mChery+) TKO mouse ovarian cancer cells grown i.p. in C57BL/6 immunocompetent mice (5,24). TKO mouse ovarian cancer cells were isolated from spontaneously formed high-grade serous ovarian cancer from a conditional Dicer-PTEN KO mouse with a p53 mutation (*p53*^LSLR172H/+^*Dicer*^flox/flox^*Pte* ^flox^/^flox^ *Amhr2*^cre/+^). Three days after inoculation, mCherry^+^ TKO cancer cells can be detected in the peritoneal cavity. This initial mCherry signal gradually dissipates 5–10 days post-injection without any treatment intervention and remains below background until about day 24 (Figure 1A). At about 25 to 30 days post-inoculation, mCherry signal returns and continues to increase, with animals developing substantial i.p. carcinomatosis and ascites by day 35 to 40 (Figure 1A-B). Thus, in this model, we observed three distinct phases of tumor progression: 1) **<u>Initial</u>** <u>phase</u> (days 1-10; yellow box); 2) **<u>Dormant</u>** <u>phase</u> (days 12-25; green) potentially associated with immune surveillance; and 3) **<u>Expansion/Immune Escape</u>** <u>phase</u> (days 25-40, blue box) due to a brake on immune surveillance (Supp Fig. 2).

**Figure 2.**
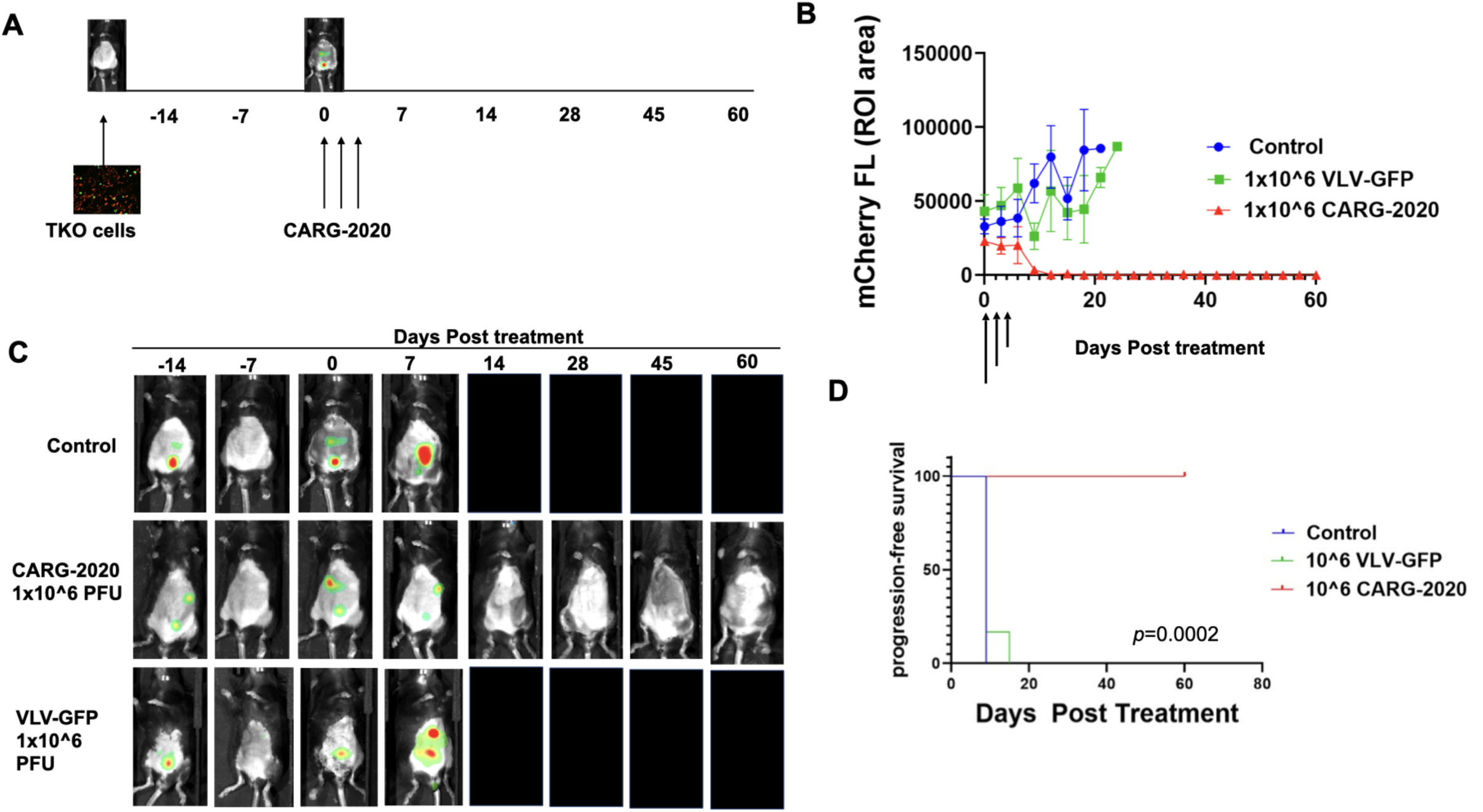
CARG-2020 causes tumor regression and significantly improves overall survival. **A.** Diagram of experimental design. 1×10^7^ mCherry+ TKO mouse ovarian cancer cells were injected i.p. in C57BL/6 mice. Mice were randomized into treatment groups and treatment began when mCherry ROI fluorescence reaches 3.2×10^8^ PFU/cell (designated as day 0). Treatment was 1×10^6^ PFU of CARG-2020 or VLV-GFP given i.p. every 3 days for a total of 3 doses (arrows show days of treatment). Control no treatment mice received PBS (n=6 per group); **B.** Tumor growth was measured by mCherry ROI fluorescence (arrows show day of treatment); **C.** Representative images showing tumor regression only in CARG-2020 treated group; **D.** overall survival was defined as day mCherry ROI area reached 1×10^9^ PFU/cell and calculated using Kaplan Meier analysis.

To determine if the same kinetics will be observed in immunocompromised mice, we injected the same number of mCherry^+^ TKO cells in athymic nude mice. Interestingly in this model, we did not observe the decrease in mCherry signal corresponding to the Dormant phase observed in C57BL/6 mice. In the athymic nude mice, the disease continuously progressed with generalized carcinomatosis by day 20-25 (Fig. 1A-B). These findings demonstrate that the presence of a functional immune system impacts the kinetics of tumor growth and suggests that it is necessary for the early control of cancer cell implantation and continued disease progression.

To test the hypothesis that the Dormant phase observed in immunocompetent C57BL/6 mice is associated with an active anti-tumor immune response, we characterized and compared the i.p. immune phenotype between naïve mice (not injected with TKO cancer cells) and mice in Dormant phase and Expansion phase using peritoneal lavage. We first looked at the CD8^+^ T cell population and observed transient but significant expansion of CD8^+^ T cells in the Dormant phase, which was lost in the Expansion phase (Fig. 1C); confirming the presence of an active immune surveillance during the Dormant phase.

Next, we evaluated the components of the innate immune response. We first focused on MDSCs since they are known to blunt protective anti-tumor immunity (37,38). Interestingly, we observed an inverse correlation between the presence of MDSCs and CD8^+^ T cells. MDSCs were not observed in the peritoneal lavage of naïve and Dormant phase mice. Instead, a significant expansion of CD11b^+^/Ly6C^+^/Ly6G^+^ granulocytic MDSC (gMDSC) was observed in the Expansion phase (Fig. 1D).

Since M2 macrophages constitute the major inflammatory component in neoplastic tissues contributing to angiogenesis and cancer metastasis, we tested for the presence of CD11b^+^/CD206^+^ M2 macrophages in the peritoneal cavity. Naïve mice were characterized by the presence of a dominant population of CD11b^low^/CD206^-^ macrophages (Fig.1E). In contrast, Dormant and Expansion phases were characterized by a dominant population of CD11b^high^/CD206^+^ M2 macrophages. The percentage of CD11b^high^/CD206^+^ M2 macrophages was significantly higher in the Expansion phase compared to naïve mice and mice in Dormant phase (Fig. 1E).

Taken together, these data demonstrate that although there is an initial anti-tumor response during the Dormant phase, the Expansion phase is characterized by the loss of CD8^+^ T cells and the expansion of immunosuppressive Ly6C^+^/Ly6G^+^ gMDSCs and CD11b^hi^/CD206^+^ M2 macrophages, which together can facilitate tumor growth and carcinomatosis (Supp. Fig. 2). We postulate that inhibiting the expansion of MDSCs and CD11b^+^/CD206^+^ M2 macrophages will aid in the promotion of CD8+ T cell expansion to consequently prevent tumor progression.

### In vitro activity of CARG-2020

To test our hypothesis that combined modulation of the innate and adaptive immune system can successfully prevent tumor progression and protect from tumor recurrence, we engineered CARG-2020, a DNA plasmid based on SFV-VSV chimeric viruses in which the native SFV glycoprotein was replaced with the VSV glycoprotein sequence (Supp. Fig. 3A). CARG-2020 carries the expression sequence for three immune modulators: IL-12, which is expected to amplify the anti-cancer response by promoting T cell proliferation; IL-17R ECD, which is expected to antagonize IL-17 signaling and block tumor-promoting inflammation; and shRNA-PD-L1, which is expected to inhibit the PD-L1/PD-1 immune checkpoint pathway (Supp. Fig. 3A).

We first tested whether CARG-2020 could infect ovarian cancer cells and express its cargo *in vitro*. Thus, TKO mouse ovarian cancer cells were exposed to CARG-2020 for 24h and IL-12 mRNA expression was quantified by qPCR. We observed a significant increase in IL-12 mRNA levels upon treatment with CARG-2020 compared to Control no treatment (Supp. Fig. 3B). Next, we determined if the mRNA expression translates into protein by quantifying secreted IL-12 in cell culture supernatants. Both the p40 and p70 IL-12 subunits were significantly higher in CARG-2020- infected cells compared to the vector Control (Supp. Fig. 3B). Western blot analysis showed protein expression of IL-12 as well as IL-17R ECD only in the CARG-2020 infected cells but not in vector Control cells (Supp. Fig. 3C). Since the action of IL-17R ECD is mediated by its binding to circulating IL-17, we examined the presence of secreted IL-17R ECD in the supernatant of cells treated with CARG-2020 by western blot analysis. As shown in Supp Fig. 3D, IL-17R ECD is detected only in supernatants of cells treated with CARG-2020 and not supernatants from cells treated with VLV-GFP control (Supp. Fig. 3D). Finally, we tested the expression of shRNA-PDL1 and it impact on PDL1 expression. Thus, cells were treated with CARG-2020 and PDL-1 expression was determined by western blot analysis in the cell lysate at 0-, 6- and 16-hours post infection. As shown in Supp. Fig. 3E, PDL-1 expression decreases in a time dependent manner in the CARG-2020 treated cells but is not affected in cells treated with vector control (Supp. Fig. 3E). These findings were confirmed by flow cytometry, which showed a decrease on PD-L1 MFI in the CARG-2020 treated cells but not with VLV-IL-12 or VLV-IL-17ECD (Supp. Fig. 3F)

**Figure 3.**
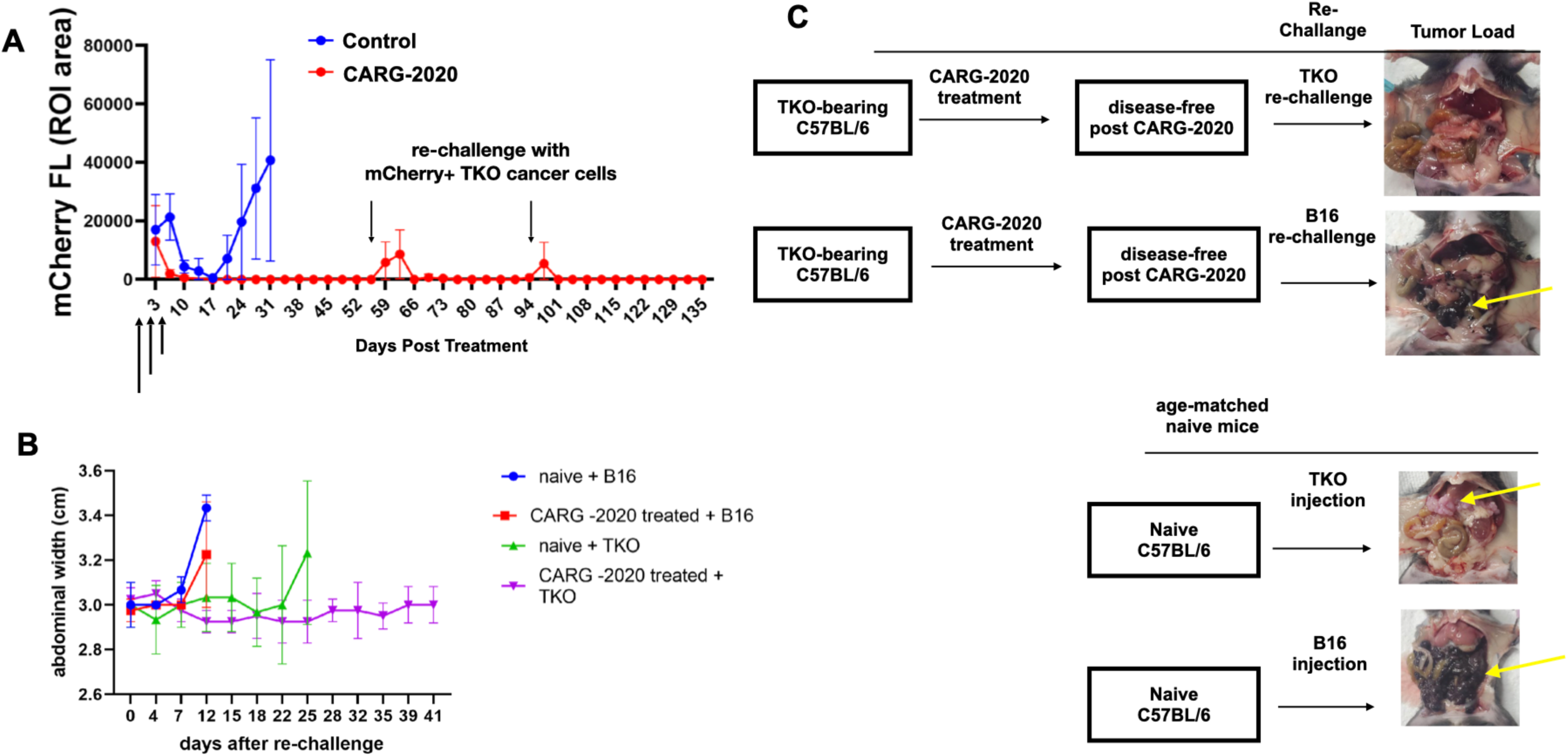
CARG-2020 provides specific immunological memory. **A.** 1×10^7^ mCherry+ TKO mouse ovarian cancer cells were injected i.p. in C57BL/6 mice. Mice were randomized into PBS control and CARG-2020 groups (n=6) when mCherry ROI fluorescence reaches 3.2×10^8^ PFU/cell (designated as day 0). Treatment was given i.p. in three doses of 1×10^6^ PFU CARG-2020 (arrows). Note disappearance of mCherry signal in CARG-2020 treatment group up to day 55, at which point mice were re-challenged with i.p. injection of 1×10^7^ mCherry+ TKO mouse ovarian cancer cells. Mice were followed for tumor formation and another re-challenge of TKO cells were given on day 94; **B.** To test for specificity, experiment in A was repeated with the addition of a group re-challenged with B16 mouse melanoma cells (n=5). Ascites formation was used as surrogate for i.p. tumor growth and monitored by measuring abdominal width. Naïve (age-matched, no tumors, no treatment) mice served as controls for i.p. growth of cancer cells; **C.** Representative necropsy images showing that CARG-2020 treatment of TKO-bearing mice protected from re-challenge with TKO ovarian cancer cells but not B16 melanoma cells. Yellow arrows point to i.p. tumors.

We next evaluated the effect the oncolytic effect CARG-2020. Thus, mCherry+ TKO mouse ovarian cancer cells were exposed to increasing concentrations of CARG-2020 or VLV-GFP and effect on cell growth and viability were monitored by real-time cell imaging. Control no treatment cells received PBS. As shown in Supplementary Figure 4A, both CARG-2020 and VLV-GFP induced a dose-dependent decrease in cell growth as measured by quantifying the mCherry region of interest (ROI) area. To determine whether the decrease in cell growth is due to cell death, we also quantified the number of cells positive for the Celltox dye (Celltox^+^). We observed a dose-dependent increase in Celltox^+^ cells in both CARG-2020- and VLV-GFP -treated cultures (Supp. Fig. 4A) compared to Control. The increase in Celltox^+^ cells peaked in the first 24 hours with CARG-2020 treatment and around 40 hours with VLV-GFP treatment (Supp. Fig. 4A). Supplementary Figure 4B shows representative mCherry and Celltox images of cultures.

To further validate that the cytotoxic effect is not limited to mice cancer cells, we treated human ovarian cancer cells, clone OCSC1-F2, with CARG-2020 and VLV-GFP. We selected the minimal cytotoxic concentration observed in the mouse cells for the human cancer and normal cells. 8 and 16 PFU/cell induce cell death in both mouse and human cancer cells but not in the normal human stromal cells (Supp. Fig. 4C). Importantly, as we recently reported, VLV vectors had no cytotoxic effect on normal human cell lines (24), and we confirmed these findings by exposing normal human endometrial stromal cells to similar doses of CARG-2020. In these cultures, CARG-2020 did not demonstrate cytotoxic activity (Supp. Fig. 4C). Together, these results demonstrate that the VLV vector is oncolytic and CARG-2020 can induce expression of its cargo *in vitro*.

To define whether the cell death observed following CARG-2020 is intrinsic to the VLV and not to the cargo, e.g. IL12; we evaluated the expression of pro-apoptotic genes, TRAIL and FAS, in ovarian cancer cells exposed to VLV-GFP, which lacks any of the immune modulatory genes. As shown in Supp Fig. 4D, exposure to VLV-GFP induces the expression of TRAIL and FAS in a time dependent manner (Supp. Fig 5A). We further validated these findings in vivo by determining TRAIL and FAS expression in tumors exposed to VLV-GFP. 24h after the administration of the second dose tumor samples were collected and examined for TRIAL and FAS mRNA expression by qPCR. We can observe a significant increase on the expression of these two apoptotic genes in tumors from animals treated with VLV-GFP (Supp. Fig 5B). These findings support our premise that the VLV component of CARG-2020 can induce apoptosis of tumor cells independently of the immune modulatory factors.

**Figure 4.**
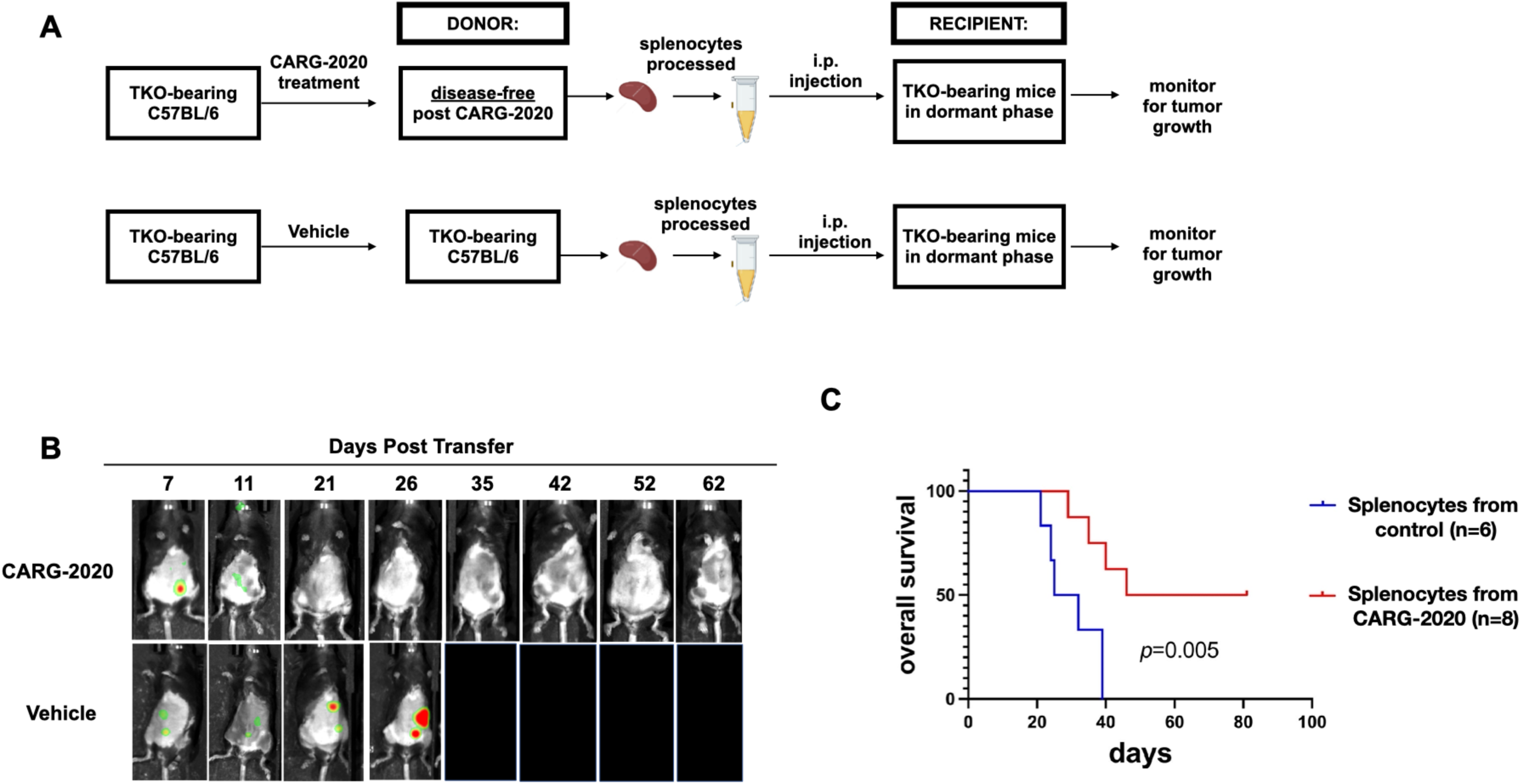
CARG-2020-induced immune memory is transferable. **A.** Diagram of spleen transfer study design. mCherry+ TKO mouse ovarian cancer cells were injected i.p. in C57BL/6 mice. Mice were treated wtih CARG-2020 (3 doses of 1×10^6^ PFU CARG-2020) groups when mCherry ROI fluorescence reaches 3.2×10^8^ PFU/sec. Splenocytes were isolated from donor mice 35 days after the start of treatment when mice were disease-free and these splenocytes were injected i.p. in another set of TKO-bearing mice in Dormant phase of the disease. Control recipients received splenocytes isolated from non-treated tumor-bearing donor mice; **B.** Representative images of Recipient mice showing mCherry+ i.p. tumor burden. **C.** overall survival was defined as day mCherry ROI area reached 1×10^9^ PFU/sec and calculated using Kaplan Meier analysis.

### In vivo efficacy of CARG-2020

Having demonstrated the *in vitro* oncolytic effect and successful expression of the immune modulatory factors carried by CARG-2020, we next evaluated it’s activity *in vivo*. Thus, we established TKO tumors i.p. in C57/BL6 mice and began treatment with CARG-2020, VLV-GFP, or PBS in the beginning of the Expansion phase as detailed in the Materials and Methods section. The time of the first dose was designated as day 0 (Fig. 2A). Three i.p. doses of 1×10^6^ PFU per dose were given every 72h and mice were imaged to follow disease progression. Mice in the CARG-2020 treated group demonstrated complete tumor regression with no detectable mCherry signal by day 14 (Fig. 2B,C). These mice remained disease-free when followed up to 60 days (Fig. 2B,C). In contrast, tumor progression and carcinomatosis was observed in mice in the VLV-GFP and PBS Control groups. Kaplan-Meier survival analysis showed significantly longer overall survival in mice treated with CARG-2020 compared to mice treated with VLV-GFP and Saline (Fig. 2D). The difference in the *in vivo* activity between CARG-2020 and VLV-GFP is interesting given that CARG-2020 and VLV-GFP are both cytotoxic *in vitro*. These findings suggest that the efficacy observed in CARG-2020-treated mice is secondary to the immune modulators carried by CARG-2020.

### CARG-2020 induces specific anti-tumor immune memory

Given that both CARG-2020 and VLV-GFP, demonstrated comparable cytotoxic activity *in vitro* (Supp. Fig. 3), we hypothesized that the superior *in vivo* activity of CARG-2020 may due to its ability to modulate the immune response. To elucidate these mechanisms, we treated a new set of mice with CARG-2020 and monitored their response. As observed previously in Fig. 2B, CARG-2020-treated mice again showed complete regression and remained disease-free for up to 55 days post-treatment (Fig. 3A). At this time, to determine the establishment of immunological memory and the capacity to prevent recurrent disease, we re-challenged these CARG-2020-treated mice with i.p. injection of mCherry^+^ TKO cells. After the re-challenge, we observed a transient mCherry signal, but this lasted for only 7 days and failed to form tumors (Fig. 3A). The mice remained disease-free for up to 5 weeks post the 1^st^ re-challenge. Thus, we re-challenged for the second time with another i.p. injection of mCherry+ TKO cells. Similarly, we observed a transient mCherry signal, but mice remained disease-free for up to 40 days after the 2^nd^ re-challenge (Fig. 3A).

To determine if this is a specific immunological memory against TKO mouse ovarian cancer cells, we established TKO i.p. tumors in another set of mice, administered three doses of CARG-2020, and re-challenged mice at day 55 as before. However, instead of re-challenging with TKO ovarian cancer cells, the mice were re-challenged with B16 mouse melanoma cells. As shown in Figures 3B and 3C, B16 melanoma cells promptly formed i.p. tumors demonstrating that previous treatment of TKO-bearing mice with CARG-2020 protects only against TKO cells. Age-matched naïve mice were used during the re-challenge experiments as control for the cancer cell growth *in vivo*. Taken together these findings demonstrate that CARG-2020 can induce a long-term and specific immune memory.

### CARG-2020-induced immune memory is transferable

To demonstrate and characterize the immunological response induced by CARG-2020, we performed splenocyte transfer (Supp. Fig. 4). mCherry+ TKO i.p. tumors were established in C57BL/6 mice followed by treatment with CARG-2020 as before. Similar to what was shown above, these treated mice demonstrated no quantifiable mCherry signal after three doses of CARG-2020 (Fig. 2B and 3A). At day 35, we collected spleens from these disease-free mice, isolated the splenocytes, and injected 5 million splenocytes i.p. into another set of TKO-bearing mice in Dormant phase of the disease (Supp. Fig. 4). In this transfer experiment, the control groups (PBS or VLV-GFP) received splenocytes from TKO-bearing mice that never received CARG-2020 and hence have measurable disease. Intriguingly, splenocytes from CARG-2020-treated mice induced tumor regression and prevented the progression of recipient mice to the Expansion phase of the disease. This translated to significant improvement in overall survival compared to Control mice (Fig. 4C). These results further demonstrate that in addition to its oncolytic activity, CARG-2020 can induce a transferable immunological memory capable of promoting an anti-tumor immune response, which is potent enough to induce complete disease regression and significantly improve overall survival.

### CARG-2020 prevents the shift towards a pro-tumor immunologic milieu and sustains a successful anti-tumor immune response

So far, our results show that CARG-2020 can promote a transferrable and specific immune memory, which is capable of inducing complete disease regression and significant improvement of survival. To fully characterize these immunological changes, we investigated the immune profile of CARG-2020-treated disease-free mice using peritoneal lavage. We observed a significant decrease in the levels of CD11b^+^/Ly6C^+^/Ly6G^+^ pro-tumor gMDSC in CARG-2020-treated mice Compared to Control (Fig. 5A). Similarly, CD11b^high^/CD206^+^ M2 pro-tumor macrophages were also significantly decreased in mice treated with CARG-2020 (Fig. 5B). In CARG-2020-treated mice, CD11b^low^/CD206^-^ macrophages were the dominant population instead (Fig. 5B). Further analysis of the macrophage population showed that CARG-2020 promoted the expansion of CD11b^+^/MHCII^+^ M1 anti-tumor macrophages (Fig. 5C).

**Figure 5.**
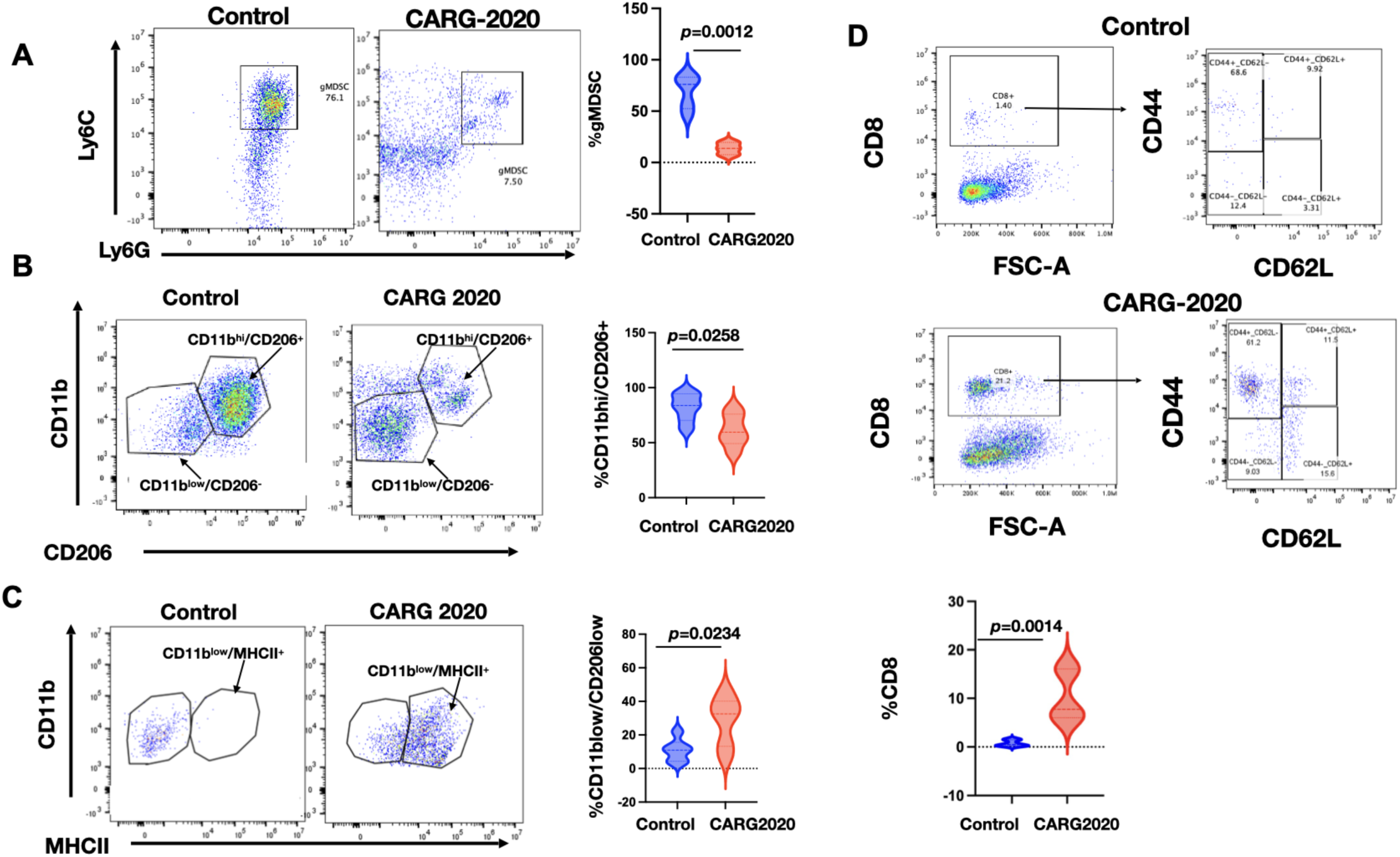
CARG-2020 modulates the innate and adaptive arms immune system. Peritoneal lavage from PBS Control and CARG-2020-treated mice (n=6) were analyzed by flow cytometry for: **A.** granulocytic myeloid suppressor cells (CD11b+/Ly6C^lo^/Ly6G+). **B.** macrophages using CD11b and CD206 and; **C.** antigen presentation using CD11b and MHCII; and **D.** CD8 memory T cells using CD8, CD44, and CD62L. Representative dot plots shown. Gating strategy is shown in Supp. Fig. 1. Data are presented as Mean ± SEM and unpaired two-tailed Student t-test was used for statistical analysis.

Finally, we characterized the adaptive immune response and determined the levels of splenic CD8^+^ T cells. We observed significantly higher percentage of CD8^+^ T cells in mice treated with CARG-2020 compared to Control (Fig. 5D). Further characterization of the expanded CD8^+^ T cell population showed that they are CD8^+^/CD44^+^/CD62L^-^ Effector Memory T cells (Fig. 5D). These data demonstrate that CARG-2020 can prevent the expansion of immunosuppressive immune cells and successfully sustain an effective cytolytic and memory T cell response.

### CARG2020 anti-tumoral effect requires adaptive immune activation

Since our data show that CARG-2020 can modulate both the innate and adaptive arms of the immune system we next determined if both are essential for CARG-2020’s anti-tumor activity. To test this hypothesis, we used athymic nude mice, which specifically lacks T cells but have an intact innate immune system. As shown above, the tumor kinetics of TKO cells in athymic nude mice do not include a Dormant phase as observed in immune competent C57BL/6 mice (Fig. 1A). We injected 5×10^6^ mCherry^+^ TKO mouse ovarian cancer cells i.p. and initiated the treatment 10 days post injection of cells. At this time, mice were randomized into Control and CARG-2020 groups. CARG-2020 was administered i.p. at 1×10^6^ PFU per dose for a total of 3 doses as shown in Figure 6A. Animal imaging showed an initial and transient decline in tumor growth in CARG-2020 treated mice but all mice eventually progressed (Fig. 6B). Flow cytometry analysis of peritoneal lavage showed no difference in the percentage of CD11b^+^/CD206^+^ M2 macrophages (Fig. 6C). However, we observed a significant increase in the percentage of CD11b^+^/MHCII^+^ M1 macrophages in mice treated with CARG-2020 (Fig. 6D, 6E). Thus, even though CARG-2020 was successful in promoting the expansion of CD11b^+^/MHCII^+^ M1 macrophages, this did not translate to an effective anti-tumor response in mice lacking T cells. These results demonstrate that modulation of both the innate and adaptive arms of the immune system is necessary for successful control of tumor growth.

**Figure 6.**
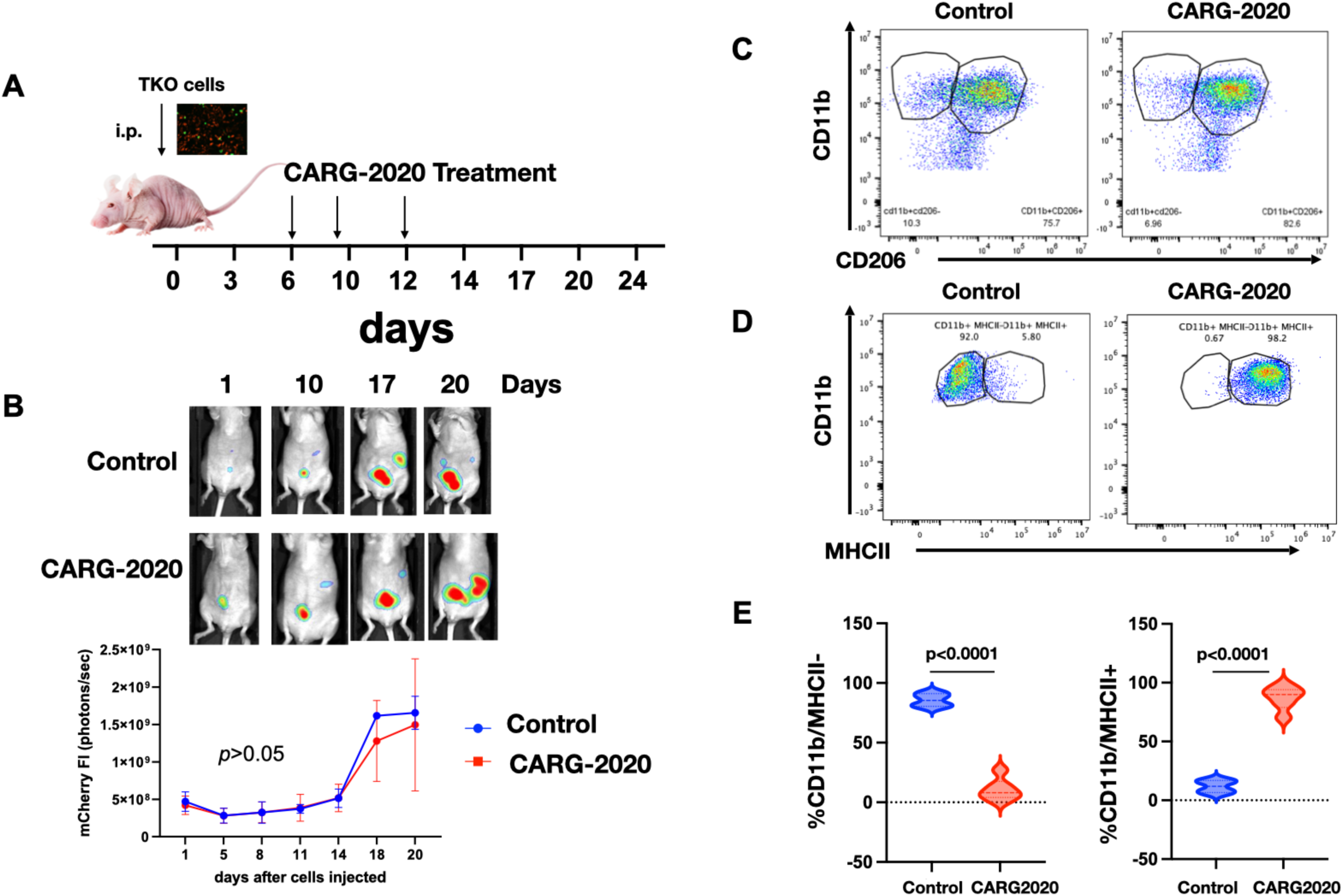
Adaptive immune system is required for CARG-2020 anti-tumor activity. **A.** Diagram of study design. TKO mouse ovarian cancer cells were injected i.p. in athymic nude mice (day 0). Mice were randomized into PBS control and CARG-2020 treatment groups on day 6 (n=5). Treatment was administered i.p. at 1×10^6^ PFU per dose for 3 doses (arrows); **B.** *top panel*, representative live animal imaging; *bottom panel*, quantification of tumor burden using mCherry ROI fluorescence area; **C.** Representative dot plots of peritoneal lavage analyzed for CD11b and CD206; **D.** Representative dot plots of peritoneal lavage analyzed for CD11b and MCHII; **E.** Quantification of D. Gating strategy is shown in Supp. Fig. 1. Data are presented as Mean ± SEM and unpaired two-tailed Student t-test was used for statistical analysis.

### Characterization of CARG-2020’s individual components

Although there is abundant information related to the anti-tumoral effect of each of the modulatory factors carried by CARG-2020, we evaluated whether the “whole is greater than its parts” by treating tumor bearing animals with VLV-IL-12, VLVIL-17DN or VLV-shRNA PDL1. We observed a strong anti-tumoral effect with VLV-IL-12, as expected, characterized by significant decrease in tumor burden (Fig. Sup. Fig. 6A). This was not the case for VLV-IL-17ECD and VLV-shRNA PDL-1 (data not shown). Although treatment with IL-12 alone represents an ideal approach for tumor immunotherapy due to its ability to activate cytotoxic T lymphocytes (39-45), it had modest effects in clinical trials (46,47). Indeed, when we compared the immunological changes induced by CARG-2020 and VLV-IL12, we observed major differences in the regulation of M1/M2 macrophages and MDSCs. CARG-2020, in addition to inhibiting tumor growth, promoted the generation of M1 macrophages and decreased the levels of CD11b^+^/Ly6C^+^/Ly6G^+^ pro-tumor gMDSC. In contrast, treatment with VLV-IL12 did not induce these immunological changes (Sup. Fig. 6B-C). These findings demonstrate that unlike CARG-2020, VLV-12 is not sufficient to modulate the innate immune response.

### Efficacy of human CARG-2020 against human ovarian cancer cells in athymic nude mice

To determine the efficacy of CARG-2020 in human cancer cells in vivo, we used a previously reported animal model consisting of mCherry OCSC-F2 human ovarian cancer cells injected i.p. into athymic nude mice (2,35,48). Animals received three injections of human-CARG-2020. Tumor growth was monitored twice a week and mCherry fluorescent ROI was plotted against time in days (Supp. Fig. 7). An initial decrease in tumor growth was observed in the CARG-2020 treated group, which was statistically significant by day 20 (p=0.0461) when compared to Control group. However, this effect was transient and tumor recurrence and growth was observed (Supp. Fig. 7A). Survival analysis did not show statistically significant differences between the groups (Supp. Fig. 7B). Taken together with the in vitro cytotoxicity data, these results demonstrate that CARG-2020 is cytolytic to human ovarian cancer cells but require adaptive immunity to provide long-term anti-tumor protection.

## Discussion

The development of intrinsic or acquired resistance is a major hurdle in the success of immune therapy in cancers, especially ovarian cancer. Resistance is typically acquired through several mechanisms targeting both the innate and adaptive arms of the immune system. Here we demonstrate that CARG-2020, an oncolytic vector containing multiple immune modulatory factors, can induce a successful anti-tumoral immune response against ovarian tumors by modifying both the innate and adaptative immune response and consequently confer a long-term protection against recurrent disease.

Dormant disseminated tumor cells (DTC) are thought to be the seed of carcinomatosis and the origin of metastatic tumors. Upon seeding, DTCs create a supportive niche that can sustain their dormant state and allow their escape from immune surveillance. These mechanisms involve decreasing cancer cell antigenicity, recruitment of MDSC, M2 macrophages, and T reg cells, and expression of checkpoints such as programmed death ligand 1 (PD-L1). Thus, a successful anti-tumor immunotherapy should have the capacity to reverse this state of tolerance and should be able to target most of the components of this complex network.

We have developed a disseminated intra-peritoneal ovarian cancer syngeneic mouse model. In this model we observed a dormant phase wherein majority of the cancer cells disappear either because they undergo cell death or are eliminated by the host immune system. However, a small number of cells not detected by imaging survive. The presence of these cells in our animal model is supported by the development of recurrent disease (expansion phase), which is observed after 10 to 15 days. The immunological milieu also changes in each of these phases. Most notable is the development of a tolerogenic condition that allows the expansion of the tumors. The main characteristics that we observed in our model during the expansion phase are: 1) suppression of CD8+ T cells; 2) recruitment of Ly6C+/Ly6G+ gMDSC and polarization of macrophages towards an M2 phenotype (Supp. Fig. 2). Our model resembles many of the findings reported in patients with ovarian cancer. It has been demonstrated *in vivo* that T-cells can halt ovarian carcinoma progression(49,50). However, antitumor immunity against established tumors is often insufficient to significantly impact tumor growth and clinical outcomes. In part, this is because ovarian cancers (and likely all solid tumors) suppress anti-tumor immunity by recruiting immunosuppressive stromal myeloid leukocytes, T regs and M2 macrophages(4,50). These cells not only have the capacity to blunt protective anti-tumor immunity (37);(50);(49); (51), but they are crucial for both the generation and maintenance of tumor vasculature as well as the promotion of metastasis (4), (52). Suppressor myeloid cells establish a pro-tumor microenvironment through multiple mechanisms including the expression of PD-L1, production of Arginase, secretion of Galectin-1 and the up-regulation of tolerogenic butyrophilins, all of which are mechanisms that can block anti-tumor immune response (4,49,53).

In this study we tested the efficacy of a novel oncolytic virus carrying immune modulatory factors targeted to “open up” the TME to T-cell control independent of tumor chemoresistance. Virus-like vesicles (VLV) are hybrid vectors based on an evolved Semliki Forest virus (SFV) RNA replicon and the envelope glycoprotein (G) from vesicular stomatitis virus (VSV) (28,30). VLVs can also express immune regulatory proteins, dominant negatives and even miRNAs and shRNA. CARG-2020 was engineered to express IL-12, antagonist to IL-17R and shRNA for PDL-1.

IL-12 has been demonstrated to regulate both innate and adaptive immunities in cancer (54). IL-12 polarizes T cells into a type 1 helper T (Th1) effector cell phenotype. Additionally, IL-12 programs effector T cells for optimal generation of effector memory T cells and T follicular helper cells (55). This cytokine was proposed as a potential agent in cancer immunotherapy due to its impressive antitumor effects in various animal models (56-58); unfortunately, the majority of clinical trials involving treatment with recombinant IL-12 failed to show sustained antitumor responses (59,60), due to the presence of a strong anti-tumoral environment and its association with toxic side effects when administrated systemically (55,61-63). We postulated that IL-12 used in combination with other immune modulatory factors and delivered within the tumor microenvironment will minimize toxicity and provide a better anti-tumor effect (64).

In this study we confirmed that IL-12 alone has anti-tumoral effect but is not able to regulate the innate immune system. IL-12 acts predominantly on lymphocytes such as T and NK cells, but its function is inhibited by factors produced in the TME, mainly by macrophages and cancer cells (64). Thus, to enhance IL-12 function, it is necessary to modulate the TME and abrogate factors that can inhibit IL-12. To achieve this objective, we introduced two additional immune modulatory factors that can reverse the status of the TME: inhibition of IL-17 and PDL-1.

IL-17, a potent proinflammatory cytokine, has been linked to inflammation and wound healing, but also it has been shown to contribute to the formation, growth, and metastasis of solid tumors (65,66). Increasing amount of evidence demonstrate that the pro-tumorigenic mechanism of IL 17 includes the promotion of an immunosuppressive endogenous microenvironment. In this regard, IL-17 works through two main mechanisms: 1) recruitment of immunosuppressive myeloid compartment either by direct secretion by TH17 cells or indirectly by cancer cells; and 2) enhancement of cancer cell survival and promotion of EMT and angiogenesis. The presence of IL-17 positive Th cells and high levels of circulating IL-17 it has been described in patients with ovarian cancer compared to healthy controls (67). Furthermore, there is strong evidences suggesting the role of IL-17 in ovarian cancer metastasis (68,69).

We reasoned that VLVs encoding the dominant-negative receptor form (dnIL-17RA) could disrupt the essential IL-17 signaling pathway. Therefore, we incorporated into VLVs, the ECD of IL-17RA but lacking the intracellular domain, as a decoy. Since the IL-17 receptor family consists of five different receptors (IL-17RA, B, C, D and E) and IL-17RA is the common subunit for all the other receptors, we generated a potent dominant-negative receptor. This mutant receptor allows the assembly of heterodimers which are competent to compete with wild-type receptors for some rate-limiting step in IL-17 signaling pathway (e.g. IL-17 ligands). In this way, the ability to induce the production of inflammatory mediators in cooperation with a wide range of ligands in the TME is abrogated and the favorable environment needed for the tumor progression is diminished.

Although immune checkpoint blockade is arguably the most effective current cancer therapy approach(70) ^4,5^, the overall response rate in most solid tumors is only around 20% (8,71). This efficacy is limited to patients with "hot" tumors, thereby warranting an effective approach to transform "cold" tumors. Oncolytic viruses are known to modulate the TME and convert cold tumors into hot tumors ^6^. CARG-2020 VLV can turn a cold tumor into hot tumor while avoiding the systemic toxicity of IL-12. Intriguingly, the heating up of the TME combined with the co-expression of the three transgenes prolonged mice survival and prevented tumor recurrence.

Conceptually the three different genetic approaches we have adopted in this study to generate CARG-2020 complement each other. First, by genetically linking p35 and p40 with a flexible linker, we have enforced co-expression of the tethered subunits, thus favoring the antitumor signaling properties of IL-12 through the JAK-STAT pathway. Second, by designing an IL-17 receptor A extracellular domain (IL-17RA-ECD) lacking the intracellular domain, the expressed IL-17RA ECD (dnIL-17RA) does not inhibit dimerization but results in a nonfunctional dimer. Thus, IL-17 downstream signaling cannot foster a favorable environment conducive for tumor progression because of lack of ability to induce the production of inflammatory mediators in cooperation with a wide range of ligands in the TME. Third, PD-L1 is expressed by diverse cell types in the TME, and high levels of PD-L1 expression dampens antitumor immunity. In addition, the PD-1/PD-L1 axis exerts a crucial role in regulating Treg development in TME. It is likely that shRNA-mediated inhibition of PD-L1 leads to the reversal of T cell exhaustion and thereby contributes to the augmentation of anti-tumor immunity. Theoretically, the suppression of PD-L1 using gene silencing approach remains by far a better option as a single interfering gene fragment is able to “switch off” protein synthesis. Since shRNA post transcriptionally inhibits PD-L1 expression, we predict that this method should prove more efficient in turning off PD-L1 expression resulting in low protein production than small molecules or antibody blockades.

Given their common roles in limiting anti-cancer immunity and promoting therapy resistance, simultaneous inhibition of IL-17 and immune checkpoints may generate synergistic anti-cancer efficacy in cancer treatment. Thus, it is plausible that each genetic component used in CARG-2020 alone is necessary but not sufficient to prevent tumor recurrence. Thus, the use of VLV as an oncolytic agent in conjunction with genetic approaches is a promising single-agent immunotherapeutic for preventing cancer recurrence.

In our model of ovarian cancer, we observed the recruitment of immunosuppressive myeloid cells during tumor expansion and hypothesized that inhibition of IL-17 could inhibit their recruitment and together with IL-12 could allow the infiltration of CD8 + cells. Indeed, treatment with CARG-2020 is associated with a significant decrease in the presence of CD11b+/Ly6C+/Ly6G+ myeloid suppressor cells and expansion of CD8+ T cells. In addition to these changes, we also observed a significant change on macrophage differentiation.

Macrophages can be phenotypically polarized by surrounding micro-environment to mount specific functional programs. M2 macrophages are a subset of what is also known as tumor associated macrophages (TAMS) and constitute a major inflammatory component in neoplastic tissues contributing to angiogenesis and cancer metastasis. They are characterized by the expression of surface markers such as CD206 and the secretion of cytokines such as IL10, CCL1, Light and IL-6.

IL-17 is one of the main regulators of macrophage polarization and is responsible for the promotion of M2 macrophage differentiation(72,73). Although it was originally considered that IL-17 was produced by the T helper cell subset TH17, numerous studies have demonstrated a wide range of innate immune cells as sources of IL-17 expression(72,74). IL-17 promotes recruitment of macrophages into the tumor microenvironment and induce their polarization towards an M2 phenotype (73,75).

In this study we focused on M2 macrophages together with MDSC since they are the more effective inhibitors CD8+ T cell activation and thus of the anti-tumoral immune response. Inhibition of both immunosuppressive cell types has been a long-time goal of immunotherapy. CARG-2020, through successful delivery of immunomodulators has successfully achieved this goal. However, we cannot ignore the potential contribution of other cell types, e.g. T regs, neutrophils, γδT cells etc. Ongoing studies are evaluating the potential contribution of these cell types in the response to CARG-2020 treatment.

Due to a high tolerogenic condition and low antigenicity, ovarian tumors are considered “cold” tumors. In order to enhance the release of antigens and reverse the “cold” status, we used the in situ vaccination approach (ISV) by promoting cell death of cancer cells together with the inhibition of M2 macrophages and MDMCs. The outcome was a specific anti-tumoral response that promotes an efficient establishment of memory, the vaccination effect, that provided a protective response to subsequent challenges (Fig. 7).

**Figure 7.**
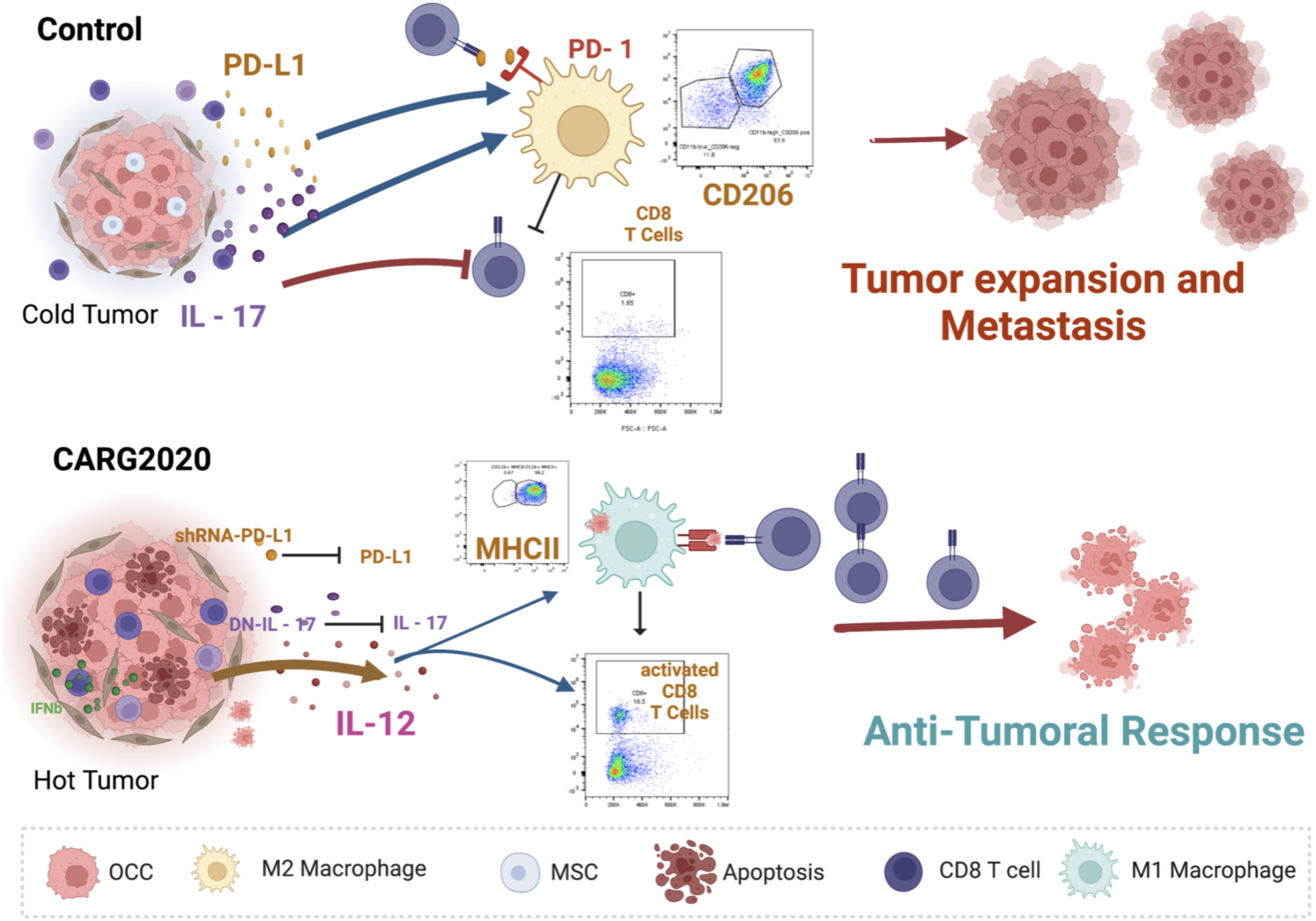
Proposed mechanism of action of CARG-2020. *top panel,* Control no treatment ovarian tumors are “cold” and express/secrete IL-17 and PD-L1 leading to differentiation of pro-tumor M2 macrophages, which inhibit the expansion of cytotoxic CD8 T cells; *bottom panel*, CARG-2020, by expressing shRNA for PD-L1 and dominant negative for IL-17, in addition to promoting the secretion of IL-12, shifts the environment to a “hot” tumor and promotes the differentiation of antigen presenting cells and consequently expansion of cytotoxic CD8 T cells.

The treatment with CARG-2020 does not induce the generation of neutralizing antibodies and is able to enhance the expression of Fas and TRAIL. Fas is a proapoptotic receptor that when expressed renders cancer cells more susceptible to its ligand (FASL), which is also expressed by cancer cells as well as by immune cells (76,77). Tumor necrosis factor (TNF)-related apoptosis-inducing ligand (TRAIL) plays an important role in apoptosis and tumor immunosurveillance (78,79). TRAIL selectively induces apoptosis in cancer cells by binding to DR4 and DR5. Increased expression of TRAIL in cancer cells has been associated with increase apoptosis in response to cellular and tissue stress (80-82).

In advance-stage disease, the tumor has established a strong immunosuppressive microenvironment (iTME) and reversing it is a challenge. Indeed, the efficacy of CARG-2020 is most optimal when given at early stages of the disease before the full establishment of iTME.

In summary, we describe a novel therapeutic approach that induce in situ vaccination by promoting cancer cells apoptosis, inhibition of MDSCs, polarization of M2 macrophages into M1 antigen presenting cells leading to an efficient anti-tumoral immune response that confers protection against potential recurrence (Fig. 11). By using the combination of well-established immune modulatory factors, we successfully demonstrate anti-tumor response and long-term protection. We postulate that the combination of oncolysis and immune modulation of the innate immune response is the most efficient way to activate a long-term protection by the adaptive immune system that will be effective in the prevention of relapse.

## Author Contributions

AA - conceptualization, methodology, formal analysis, supervision, visualization, writing

AF – investigation, validation, formal analysis

BM – investigation

MK – investigation

HC – investigation

RG – conceptualization

VN – conceptualization

BJ – conceptualization

TY – conceptualization

TR - conceptualization

GM - conceptualization, methodology, formal analysis, visualization, writing, funding acquisition

## Acknowledgment

This research was funded in part by the Janet Burros Memorial Foundation; the National Institute of Health, National Institute of Diabetes and Digestive and Kidney award (NIH NIDDK R44DK113858) and National Institute of Allergy and Infectious Diseases award (NIH NIAID R43AI149798) to V.N.

