## Supplementary Figures for "Immune modulation of innate and adaptive responses restores immune surveillance and establishes anti-tumor immunological memory"

Supplementary Figure 1

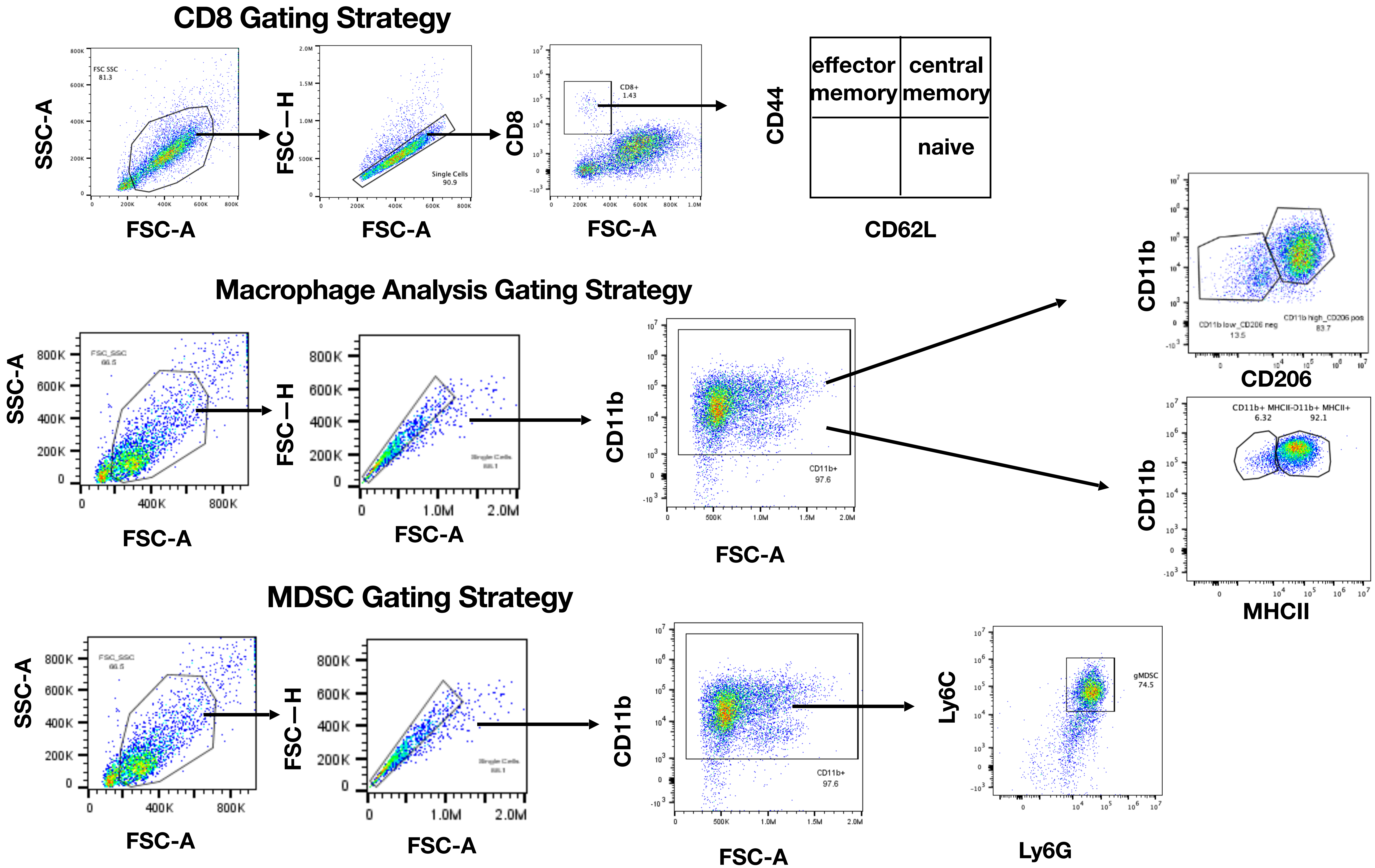

#### Supplementary Figure 2

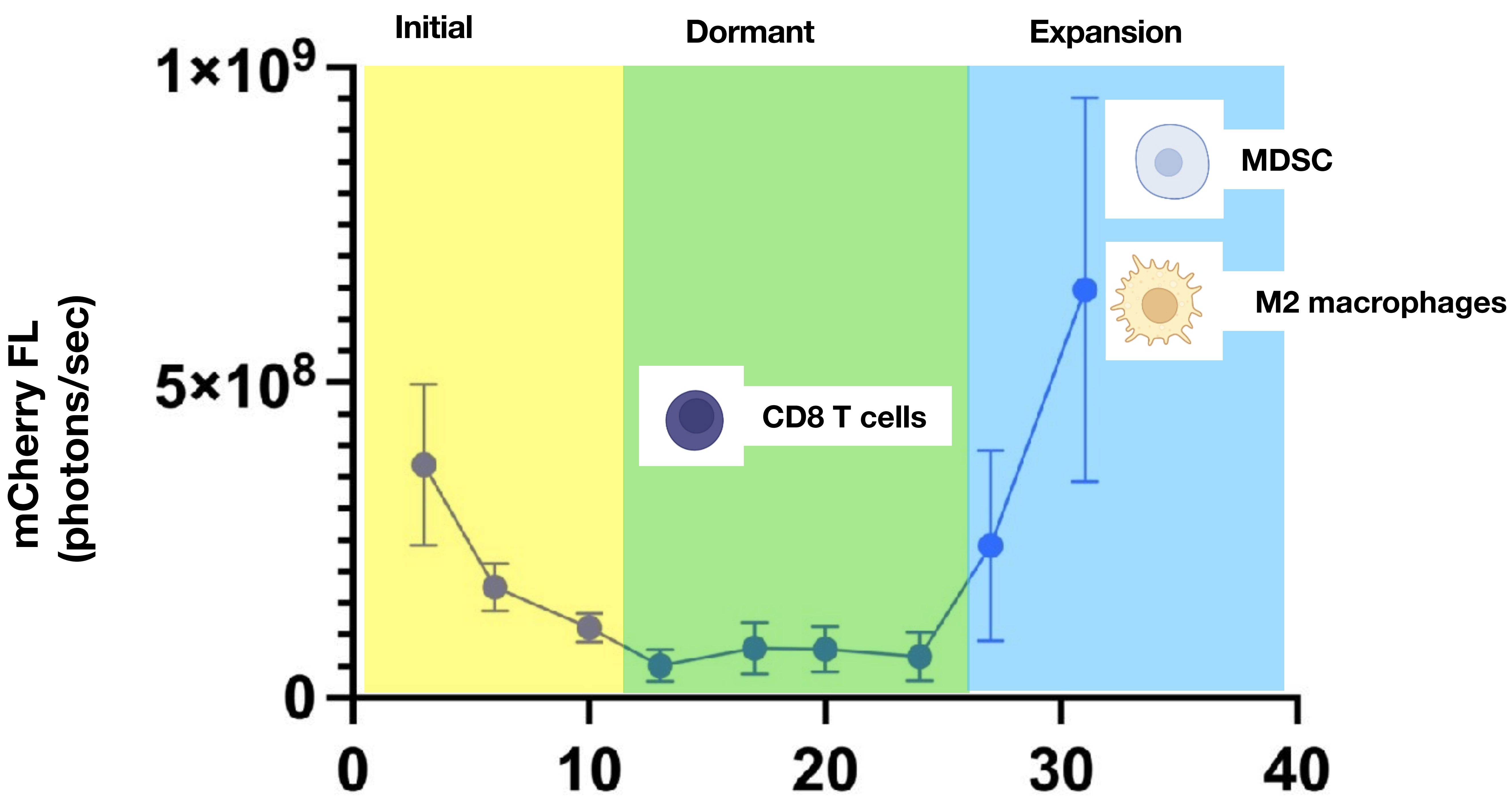

### Supplementary Figure 3

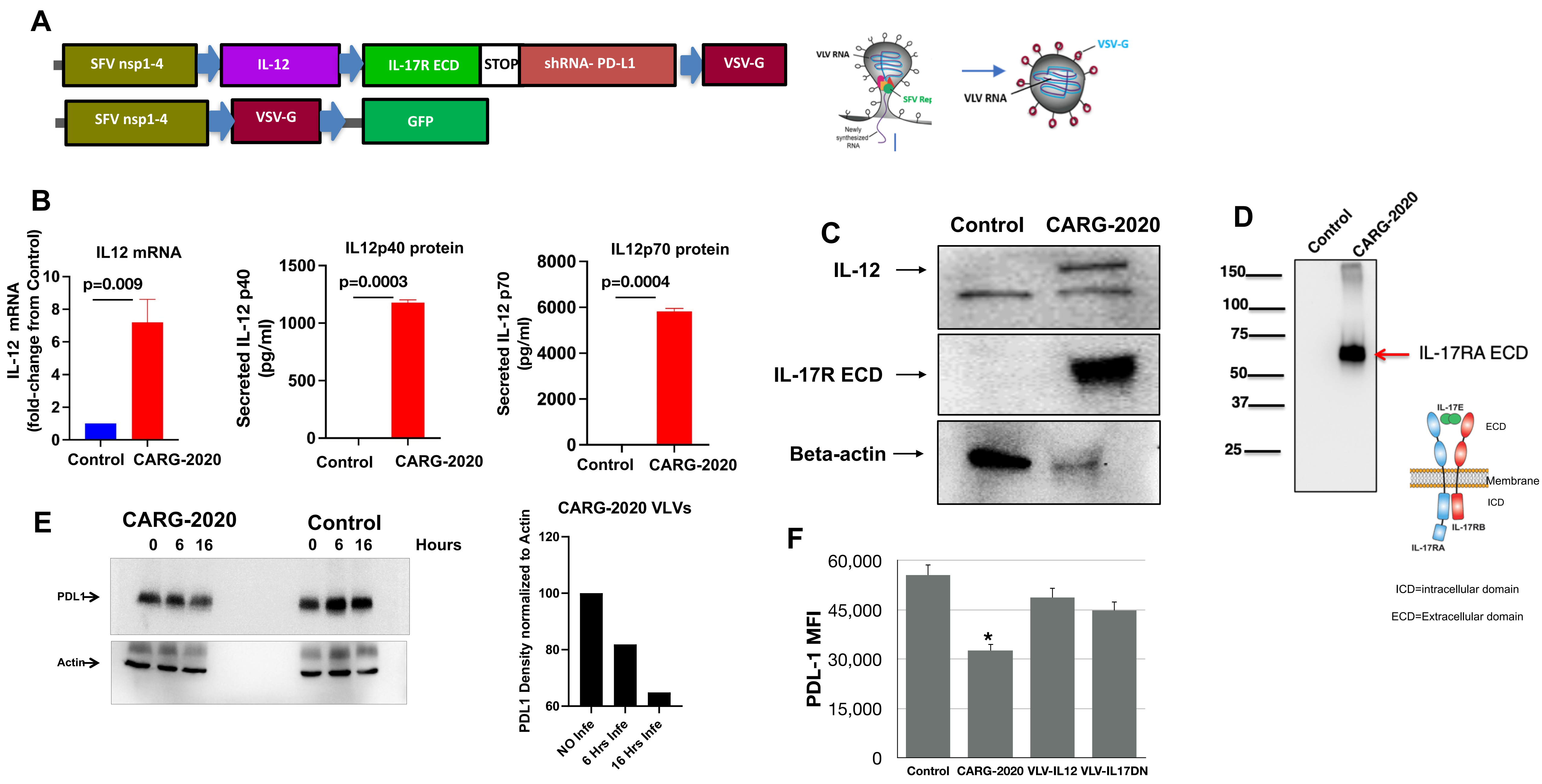

#### Supplementary Figure 4

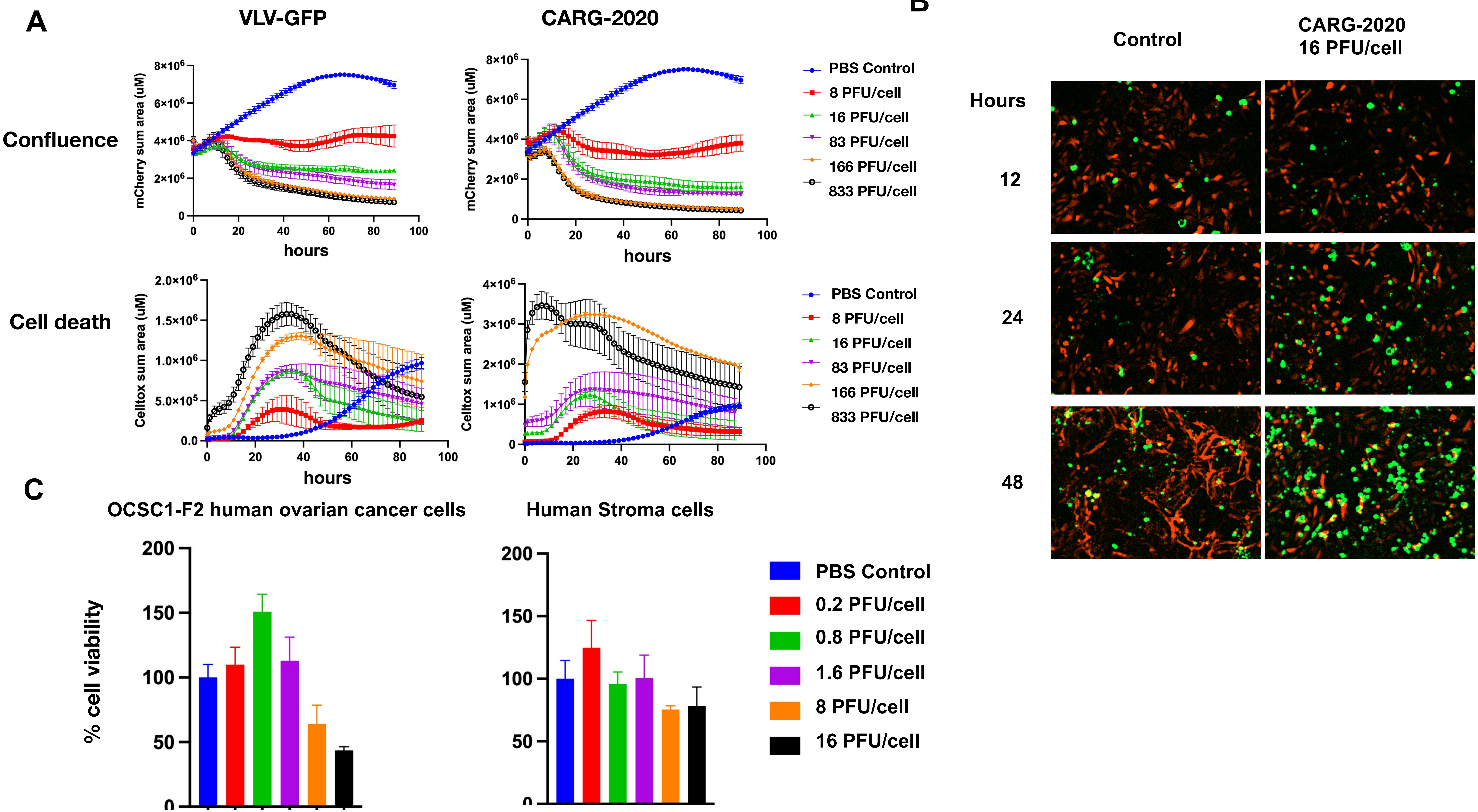

#### Supplementary Figure 5

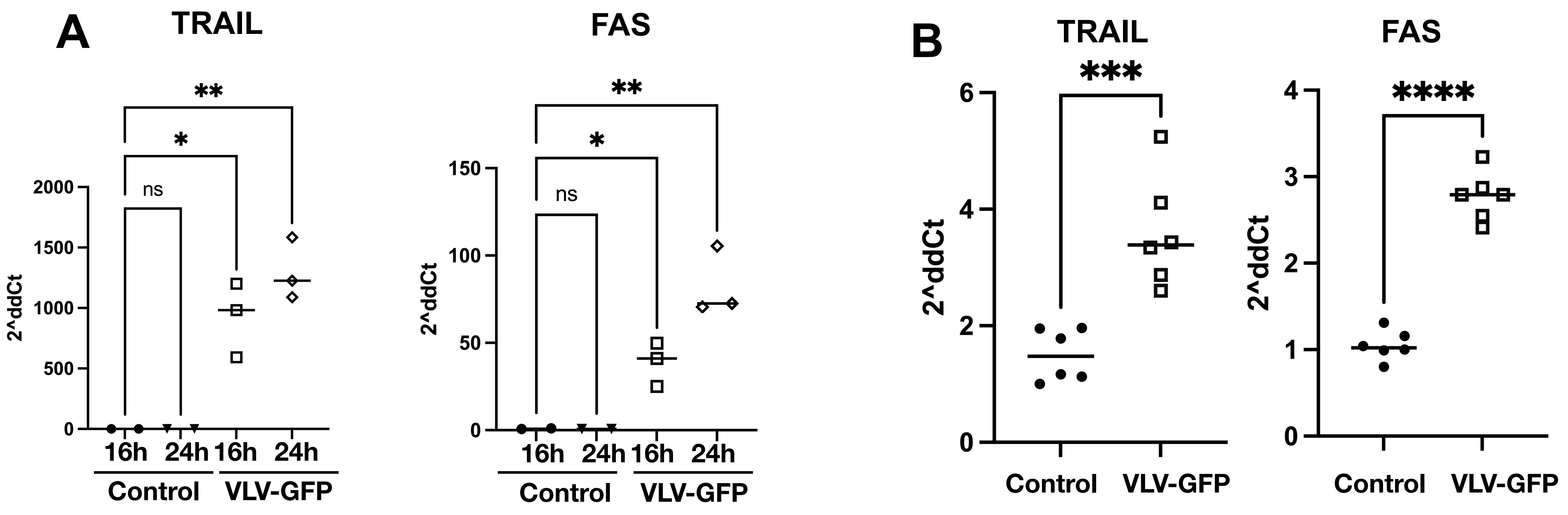

#### Supplementary Figure 6

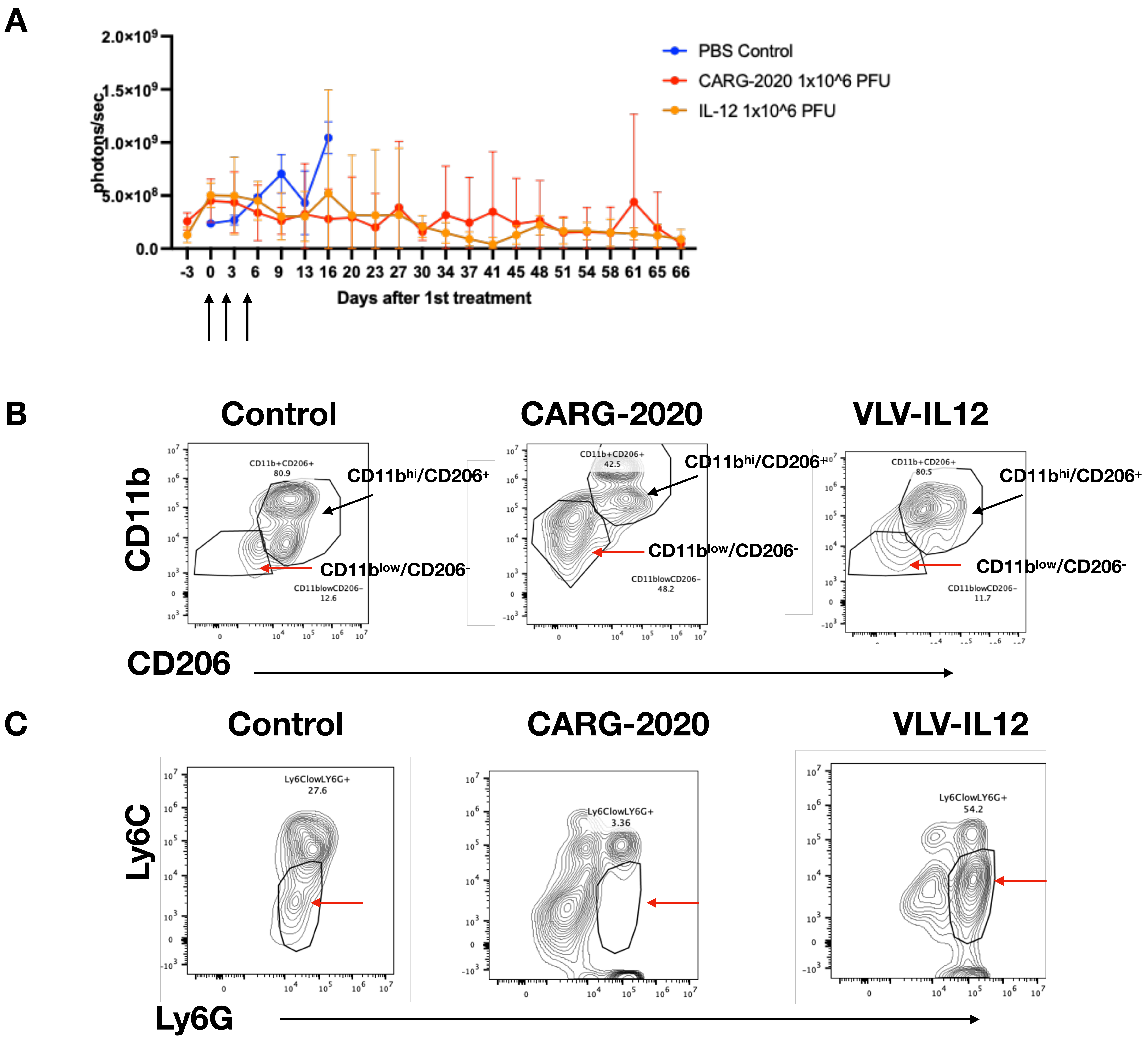

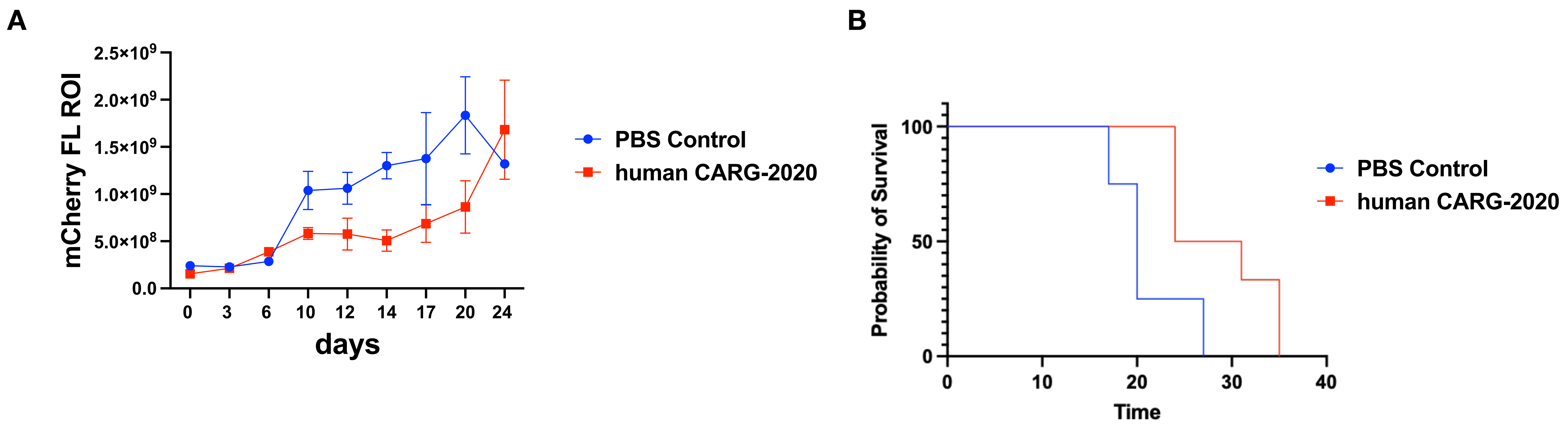
A
mCherry FL ROI
2.5×10^9
2.0×10^9
1.5×10^9
1.0×10^9
5.0×10^8
0.0
0
3
6
10
12
14
17
20
24
days
PBS Control
human CARG-2020
B
Probability of Survival
100
50
0
0
10
20
30
40
Time
PBS Control
human CARG-2020
