## Supplementary Figure Legend for "Immune modulation of innate and adaptive responses restores immune surveillance and establishes anti-tumor immunological memory"

**Supplementary Figure 1**

Flow cytometry gating strategy CD8 T cells, Macrophages and MDSC.

**Supplementary Figure 2**

Diagram showing major immune phenotypes in each phase of ovarian tumor progression.

**Supplementary Figure 3**

**A.** Diagram of CARG-2020 vector and GFP control vector; **B.** TKO mouse ovarian cancer cells were treated *in vitro* with PBS Control or $1x10^5$ PFU/cell of CARG-2020 for 24h. IL12 mRNA was quantified by qPCR and secreted IL-12p40 and p70 protein were quantified using xMAP technology; **C.** Expression of IL-12 and IL-17R ECD were detected by western blot analysis in total cell lysate. **D.** Secretion of IL-17 RA-ECD was detected by western blot analysis in the supernatant of cells treated with CARG-2020. Control cells were treated with VLV-GFP vector. Representative imagine of 3 independent experiments.

E. Expression of PDL-1 following treatment with CARG-2020. Note the time dependent decrease on PDL-1 expression on cells treated with CARG-2020. Control=VLV-GFP. Representative imagine of 3 independent experiments.

**Supplementary Figure 4**

**A.** TKO mouse ovarian cancer cells were treated with increasing concentrations of CARG-2020 or VLV-GFP and effect on cell growth and cell death is quantified in real time using Cytation5/Biospa by measuring mCherry confluence and Celltox$^{TM}$ fluorescence; **B.** Representative fluorescence images showing PBS Control and CARG-2020-treated cultures. *red*, mCherry; *green*, Celltox$^{TM}$; **C.** Human ovarian cancer cells, clone OCSC1-F2 and the hTERT-immortalized human endometrial stromal cells were treated with increasing concentrations of CARG-2020 for 72h and cell death was quantified using Celltox$^{TM}$ fluorescence. Note that CARG-2020 induces cell death only in cancer cells and not in normal cells.

**Supplementary Figure 5**

**A**. Cytolytic effect of VLV-GFP in cancer cells is mediated by the induction of pro-apoptotic genes. Treatment of cancer cells with VLV-GFP is associated with a significant increase on TRAIL and FAS mRNA expression. *=$p>0.005$; **= $p>0.001$

**B.** In vivo induction of pro-apoptotic genes. Mice bearing mCherry-TKO tumors were treated 2X with VLV-GFP. The expression of TRAIL and FAS was determined 24h after the last treatment. Note the significant increase on TRAIL and FAS mRNA expression following VLV-GFP treatment. ***= $p>0.0001$

**Supplementary Figure 6**

**A**. Antitumoral effect of VLV-IL12 and CARG-2020 in mouse ovarian cancer. Tumor growth was measured by mCherry ROI fluorescence (arrows show day of treatment).

**B**. CARG-2020 but not VLV-IL-12 modulates the innate immune system. Peritoneal lavage from VLV-IL-12 and CARG-2020-treated mice (n=6) were analyzed by flow cytometry for macrophages using CD11b and CD206. VLV-IL-12 treatment is not able to increase the number of $CD11b^{low}/CD206^{-}$ macrophages compared to CARG2020. Representative dot plots shown. Gating strategy is shown in Supp. Fig. 1.

**C.** Decrease on Ly6C+/Ly6G+ MSC is observed only in the CARG2020 treated group but not in those treated with VLV-IL-12 group. Representative dot plots shown. Gating strategy is shown in Supp. Fig. 1.

**Supplementary Figure 7**

**A**. Mean tumor burden over time of nude mice bearing OCSC1-F2 human ovarian tumors. mCherry fluorescent ROI was used to provide a measure of tumor burden. Data are presented as mean ± SEM. Treatments began on day 6 and continued through to day 12.

**B**. Survival analysis using Kaplan-Meier.
